# Highly diverged non-vertebrate Ki-67 proteins with conserved biophysical and functional properties

**DOI:** 10.64898/2026.06.18.733134

**Authors:** Austin Haider, Aleksandra Zieminska, Geronimo Dubra, Kari Gaalswyk, Lili Houston, Michael Phillips, Stéphane Guindon, David Llères, Kingshuk Ghosh, Daniel Fisher, Liliana Krasinska

## Abstract

The Ki-67 protein is a widely used marker of mammalian cell proliferation that plays critical roles in heterochromatin organisation and mitotic perichromosomal layer formation. Thus, its apparent restriction to vertebrates is puzzling. However, sequences with limited homology to subdomains of Ki-67 are present in genomes from some non-vertebrate species. Here, using structural modelling, biophysical and functional analysis, we identify Ki-67-like proteins across diverse non-vertebrate eukaryotes, including tunicates, sponges, molluscs, insects and fungi. Despite extremely low sequence homology, the disordered domains of Ki-67 homologues have generally conserved biophysical and molecular properties. This is further reflected by comparison of cellular localisation, dynamics and effects on chromatin organisation of human, *Ciona* and *Drosophila* Ki-67 homologues. These results reinforce the emerging paradigm that functions may be encoded by general biophysical features in highly diverged intrinsically disordered domains. Our approach provides a blueprint for comparative biology of disordered proteins and a foundation for future studies exploring functions of Ki-67 homologues across eukaryotes.

## Introduction

The cell proliferation marker Ki-67 is a large (> 300kDa in mice and humans) nuclear protein expressed in all proliferating mammalian cells. Though Ki-67 expression is cell cycle-regulated, it is not essential for cell division. It is a hub for protein-protein interactions ^1^, a property that probably contributes to its mitotic function as a platform for the formation of the perichromosomal layer (PCL) ^1–3^. Ki-67 is essential for recruitment of other nucleolar components to this compartment around condensed chromosomes, allowing their equal partitioning into daughter cells ^1–3^. In interphase, Ki-67 associates with constitutive heterochromatin, which is found in pericentromeric and telomeric regions of chromosomes and is characterised by the histone marks H3K9me3 and H4K20me3. Disruption of Ki-67 expression causes a general decrease in chromatin compaction, including perinucleolar and pericentromeric heterochromatic regions ^1,4^. This results in widespread deregulation of the entire transcriptome, reducing cell plasticity, and affecting all stages of tumourigenesis ^5^.

Despite these fundamental roles, the biochemical properties of Ki-67 are poorly understood. This is in part because the protein has no enzymatic activity and most of its sequence is intrinsically disordered. The only fully structured region is the N-terminal forkhead-associated (FHA) domain of ≈100 amino acids. FHA domains are phospho-threonine binding modules found in all eukaryotes. In humans alone, >100 proteins with diverse functions contain FHA domains, and, as such, the FHA domain gives few clues to the functions of Ki-67. The human Ki-67 FHA domain interacts with NIFK ^6,7^, a nucleolar protein present in all eukaryotes that is essential in human cells, as well as in budding and fission yeasts ^8,9^, but the role of this interaction is not known. A large part of Ki-67 is formed by repeats, whose function is also unknown. Human Ki-67 contains 16 such repeats of 122 amino acids with 40-60% identity, while other vertebrate Ki-67 homologues contain a variable number of repeats of different lengths and lower sequence conservation (for example, *Xenopus laevis* Ki-67 has 13 repeats 29 aa long).

Ki-67 homologues have not been described outside of vertebrates (see ^10^). Several non-exclusive hypotheses could be invoked to explain why simpler eukaryotes might not need Ki-67. Vertebrates might require Ki-67 to compartmentalise their larger genome (3200 Mb in human, compared with 165 Mb in *Drosophila melanogaster* or 14 Mb in *Schizosaccharomyces pombe*). Fungi often have a closed mitosis, where nucleoplasm does not mix with the cytoplasm, and as such might not require a PCL-based mechanism for dividing nucleolar components. However, a simpler explanation might be that the high evolution rate of intrinsically disordered regions (IDRs) has hindered the discovery of invertebrate Ki-67 homologues. The conservation of heterochromatin throughout eukaryotes and the intrinsically disordered nature of Ki-67 led us to hypothesise that Ki-67 homologues might exist in other branches of the tree of life, and that conservation of molecular features encoded in the sequence, which frequently occurs in intrinsically disordered proteins (IDPs) ^11,12^, might nevertheless allow their identification. By applying structural modelling, biophysical analysis and functional experiments in living cells, we identify Ki-67 homologues in a wide range of non-vertebrate eukaryotes.

## Results

### Identification of ascidian Ki-67 homologue

We first asked whether Ki-67 exists in tunicates, invertebrate chordates whose ancestors diverged from vertebrates at the root of the phylogenetic tree. A TBLASTN (ClusteredRN) search against the commonly used tunicate model *Ciona robusta* (formerly known as *Ciona intestinalis*) sequences with human Ki-67 returned two proteins, of 1552 amino acids (aa) (XP_002132161.1) and of 1418 aa (XP_018673001), while an unbiased query using the more sensitive phylogeny-based SHOOT ^13^ against all species revealed two *Ciona* UniProt entries: a 267 aa long FHA domain-containing protein (F6VDD5) and a 162 aa uncharacterised protein (H2XQ73). The F6VDD5 protein is encoded on chromosome 2, adjacent to a 270 aa uncharacterised protein (F6UXW5; position 7.250.602-7.253.841), with the H2XQ73-encoding gene located immediately downstream (7.254.208-7.258.365). BLAST search with *Ciona* F6VDD5 protein against human revealed as top hit Ki-67, with 50% identity within the first 100 residues, which correspond to the FHA domain (Fig. 1A). The F6UXW5 *Ciona* protein shares 23% identity with the 648-776 region of human Ki-67, while H2XQ73 can be aligned to the repeat domain of human Ki-67, with 30% identity with residues 1312-1453, and lower values for downstream repeat domain fragments. We surmised that the three genes present in UniProt are misannotated and in fact correspond to a single gene encoding a putative *Ciona* Ki-67 protein with limited homology to human Ki-67 outside of the FHA domain. The ascidian genomic Aniseed database (https://www.aniseed.fr) revealed several genes including one in *Ciona robusta* (KH2012:KH.C2.974.v2.A.nonSL1-1), that maps to the same position on chromosome 2 (7.250.605-7.261.095; Fig. 1B) as the three genes annotated in UniProt. Homologues are also present in other ascidians, *Phallusia mammillata*, *Phallusia fumigata*, *Molgula oculata* and *Molgula occidentalis* (Supp. Fig. 1A). The Ciona gene encodes a 1571 residue protein, with Pfam-predicted N-terminal FHA and protein phosphatase 1 (PP1)-binding domains. Alignment of the Aniseed gene with the BLAST hits suggests an incomplete annotation of the 1552 aa protein (XP_002132161.1), that lacks residues 537-555, while the 1418 aa protein is also missing the N-terminus. In summary, our analysis revealed the existence of a single Ki-67 homologue in Ciona.

**Figure 1.**
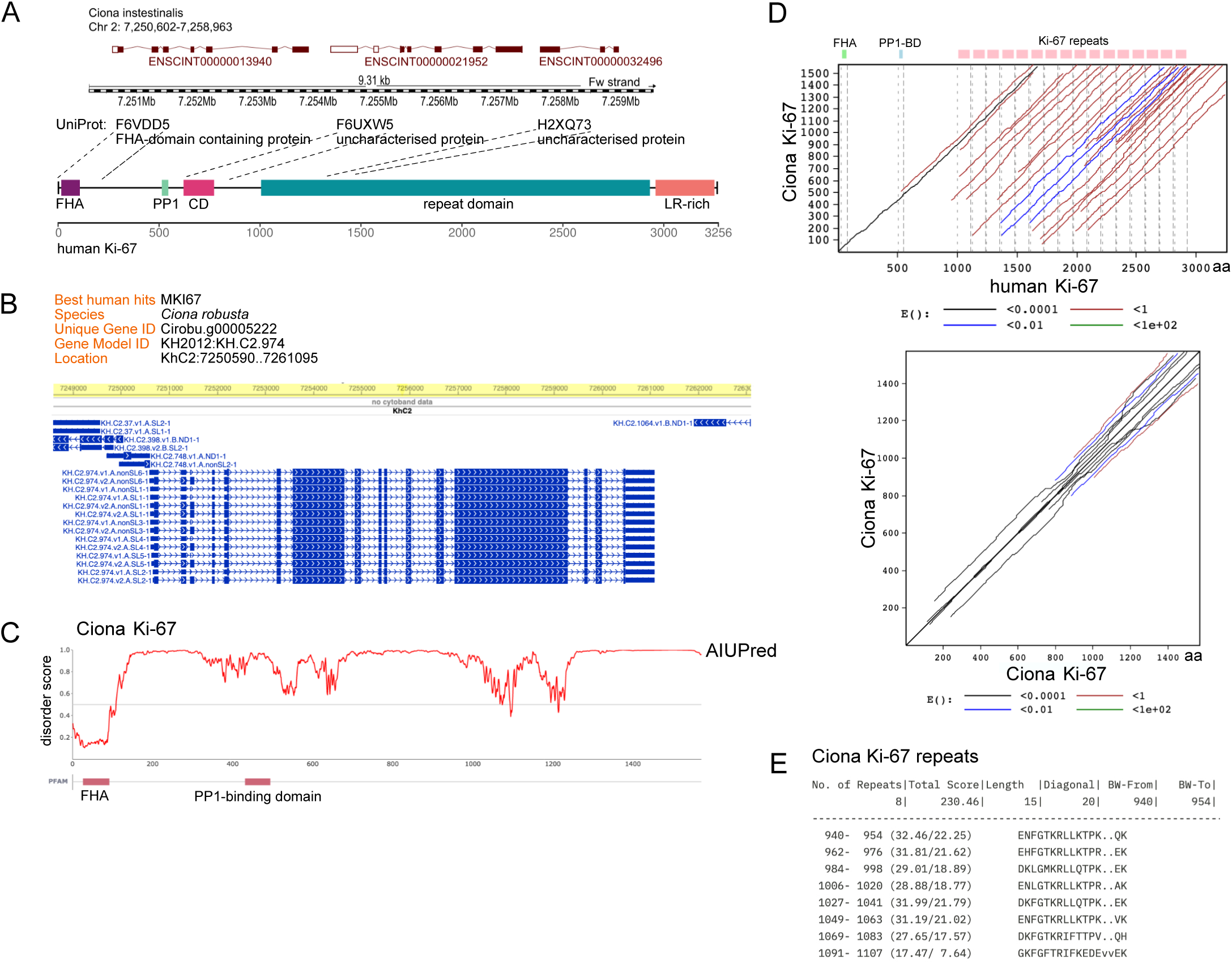
Identification and characterisation of Ciona Ki-67 homologue. A. Schematic presentation of UniProt Ciona Ki-67 entries encoded by three genes on Chr. 2 (Ensembl genome browser) and mapped to human Ki-67 protein. B. The *Ciona robusta* Chr. 2 region encoding the three genes (A) annotated in Ensemble corresponds to one protein-coding gene according to Aniseed database. C. AIUpred disorder prediction plot of the Ciona Ki-67 homologue. D. Top, LALIGN plot aligning human and Ciona Ki-67 proteins, showing similarity scores; bottom, PLALIGN dot plot-like graph identifying internal duplications wihin Ciona Ki-67 protein. E. Repeats identified in Ciona Ki-67 by RADAR.

AIUPred3 ^14^ predicts that, like vertebrate Ki-67, the *Ciona* protein is intrinsically disordered outside of the FHA domain (Fig. 1C); it has a PP1-binding motif in its N-terminal part (Table 1). BLAST query of the *Ciona* genome with human Ki-67 omitting these two features failed to return any hits, indicating insufficient homology in the disordered domain. Nevertheless, pairwise local comparison with human Ki-67 using the internal duplication finder PLALIGN ^15^ showed statistically significant alignment throughout the entire length of the *Ciona* protein (Fig. 1D, top), while self-alignment indicated duplications within its sequence (Fig. 1D, bottom). Detection of repeats using the RADAR analysis tool ^16^ revealed a 15 aa sequence repeated 8 times within the 940-1107 aa region (Fig. 1E). Like human Ki-67, the *Ciona* protein has a highly basic isoelectric point (10.19 and 9.82, respectively). Furthermore, its expression profile suggests that, like mammalian Ki-67, the Ciona protein is expressed in proliferating cells: from unfertilised egg to stage 26 (hatching larva), the mRNA profile is similar to that of the *Ciona* homologue of Cyclin A, a cell cycle regulator, and is clearly distinct from that of the transcription factor TBXT, required for mesoderm formation and differentiation (Supp. Fig. 1B). Thus, despite a limited overall homology, the *Ciona* protein shares a number of general features with human Ki-67: it is a basic protein that is intrinsically disordered with the exception of an FHA domain at its N-terminus, it contains a PP1-binding domain, several repeats, and multiple CDK phosphorylation motifs; and it is expressed in proliferating cells.

### Ciona Ki-67 FHA domain sequence and interaction are conserved

Given the higher sequence conservation of structured domains, we surmised that any FHA domain features that are specific to Ki-67 and conserved between vertebrates and *Ciona* might enable to identify other invertebrate Ki-67 homologues. The sequence homology of FHA domains of different human proteins (113 in total, according to the Small Modular Architecture Research Tool, SMART; http://smart.embl-heidelberg.de/) is low (Fig. 2A). The few conserved residues (GRSHNN) are involved in binding phospho-threonine and surrounding amino acids. Yet all FHA domains present a very similar three-dimensional structure, where 11 β-strands form two β-sheets that position the loops which interact with phospho-threonine residues (Fig. 2B). We suspected that Ki-67 FHA domain sequences are more highly conserved. Indeed, the InterPro database ^17^, that classifies proteins into families, integrating various protein signature databases into one resource, has a separate entry for Ki-67-like FHA domains. Using this resource and SHOOT, we retreived a selection of vertebrate Ki-67 FHA domain proteins, including evolutionary distant organisms (*e.g.* ghost shark, hereafter “shark”) and the largest living mammal, the blue whale (hereafter “whale”). Alignment of FHA domains of various vertebrate Ki-67 homologues and that of *Ciona robusta* (Fig. 2C) showed markedly higher sequence identity than unrelated FHA domains (Fig. 2A), indicating unusually high conservation within the Ki-67 FHA family. The AlphaFold3-predicted 3D structure of *Ciona* Ki-67 FHA domain was essentially identical to that of human Ki-67 (Fig. 2B). The structure of human Ki-67 FHA-NIFK interaction has been solved by NMR, revealing a unique feature, where the interaction between the two proteins is extended beyond the residues surrounding phospho-threonine and involves a β-sheet formed by NIFK that stacks with the β4 strand of the FHA ^7^ (see Fig. 2E). We speculated that this unusual mode of interaction might be conserved in Ki-67 and NIFK homologues. The alignment of FHA domains of vertebrate Ki-67 homologues showed that the residues interacting with NIFK ^7^ are strictly conserved between different species (Fig. 2D, with logo), but they are absent in sequences of FHA domains of other proteins (see Fig. 2A). AlphaFold3 ^18^ correctly predicted the structure of human Ki-67 FHA domain binding the NIFK C-terminal peptide with phosphorylated threonines (Fig. 2E). The residues forming the β-strand anti-parallel to the β4 strand of the FHA domain are conserved between human (DDEIVFK) and *Ciona* (DEEVVFK), but the threonines (T234 and T238) that interact with the FHA domain loops of human Ki-67 are not conserved in *Ciona* NIFK (Supp. Fig. 1C). Interestingly, in *Ciona* NIFK, CDK-phosphorylatable threonines (followed by a proline) are located C-terminally: T247, directly after the anti-parallel β-strand, and T274, that interacts with the FHA domain loops (Fig. 2E). This suggests that the formation of the anti-parallel β-strand, which strengthens the interaction between NIFK and the FHA domain, is an evolutionarily early and conserved feature, and that there are additional sites of the FHA domain that can bind phospho-threonines outside of the loops. We also modeled the interaction between the *Ciona* Ki-67 FHA domain with the C-terminal fragment of human NIFK, phosphorylated on threonines. Interestingly, the binding was identical to the binding of human Ki-67 to NIFK (Supp. Fig. 1D), indicating strong conservation of structural properties. Altogether, these results strongly suggest that the protein identified in *Ciona robusta* is a Ki-67 homologue (we will refer to it as CiKi-67 thereafter).

**Figure 2.**
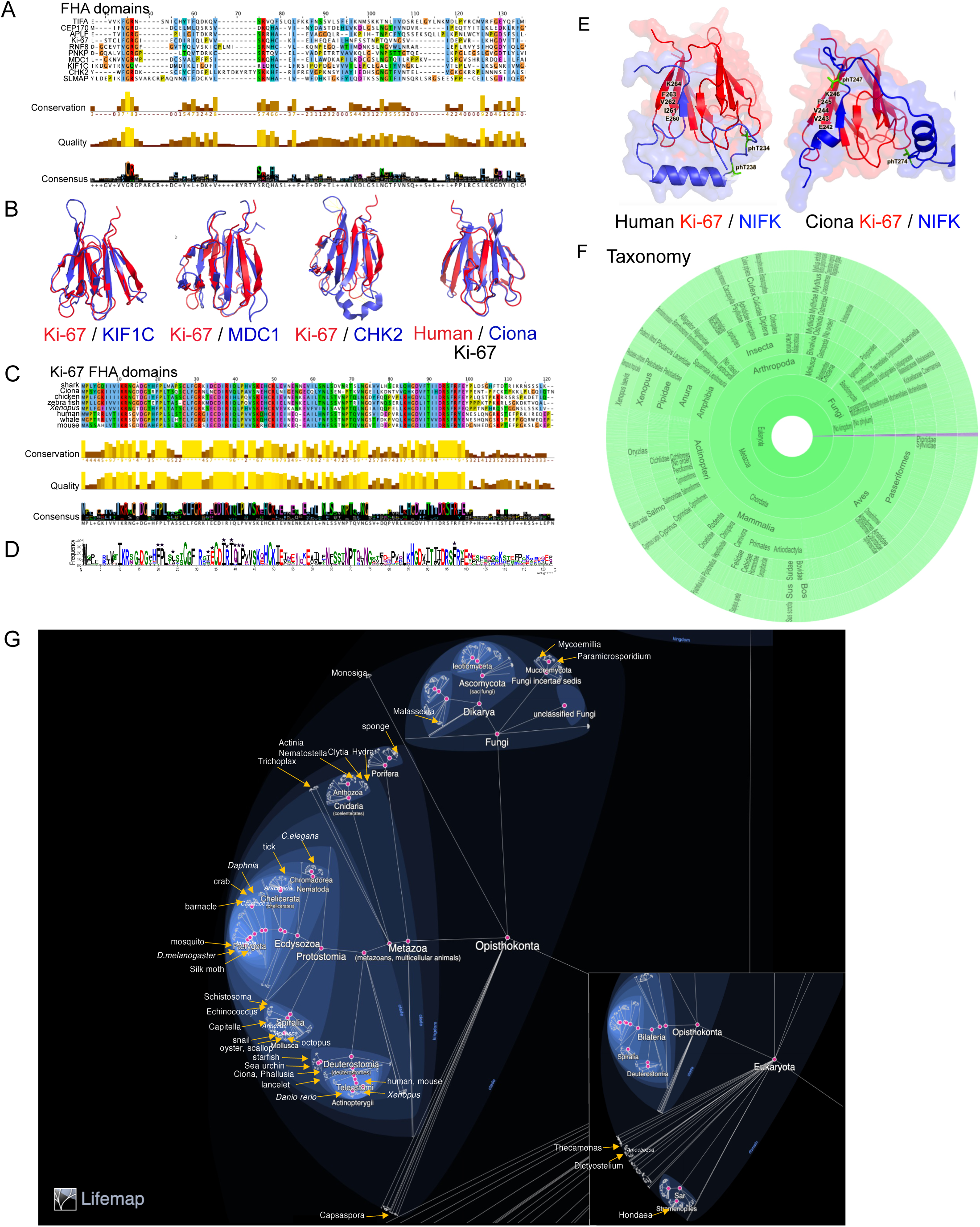
Conservation of Ki-67 FHA domain sequence and interaction with NIFK. A. MUSCLE (MUltiple Sequence Comparison by Log-Expectation) alignment of FHA domain sequences of the indicated human proteins, coloured in Clustal. B. Alignment of 3D structures of FHA domains of human Ki-67 (PDB 1R21) with human KIF1C (PDB 2G1L), MDC1 (PDB 3VA4), and CHK2 (PDB 1GXC); and with Ciona Ki-67 (for Ciona, AlphaFold3 model prediction was used, aa 1-100). Images were generated using PyMol. C. MUSCLE alingment of FHA sequences of vertebrate and Ciona Ki-67 homologues, coloured in Clustal. D. Consensus logo of vertebrate and Ciona Ki-67 FHA domain sequences; asterics indicate residues interacting with NIFK. E. Comparison of human and Ciona Ki-67 FHA domain interaction with NIFK (Ciona NIFK: UniParc entry XP_002128246.1, aa 234-297) predicted by AlphaFold3. NIFK residues forming the antiparalel B-strand and phosphorylated Thr are indicated. Images were generated using PyMol. F. Sunburst view of over 3k taxa identified by InterPro database as FHA domain-containing Ki-67 and similar proteins. G. Lifemap illustration of species whose Ki-67 homologues were analysed in this study.

### Identification of other non-vertebrate Ki-67 homologues

We next tested whether this striking FHA domain sequence and structure conservation can identify other non-vertebrate Ki-67 homologues. The InterPro Ki-67 FHA entry contains over 2000 eukaryotic proteins, most of which belong to vertebrate species. However, there are proteins listed from multiple non-vertebrate species, and even some bacteria (Fig. 2F). We asked whether the latter are genuine Ki-67 homologues. For this analysis, we chose representantatives of various classes of vertebrates, including model organisms (mouse, *Xenopus laevis* and *Danio rerio*). We then selected a wide range of invertebrate animal phyla and classes (Echinodermata, Mollusca, Annelida, Platyhelminthes, Insecta, Crustacea, Arachnida, Nematoda, Cnidaria, and Porifera), organisms from outside Metazoa, such as fungi, and non-Opisthokints (Fig. 2G; ^19^). Among the non-Metazoa, we considered putative homologues from the following species: i) *Trichoplax adhaerens*, together with three other species, forms the phylum Placozoa, a basal group of multicellular blob-like animals; ii) *Capsaspora owczazaki*, an ameboid protist, is one of the closest unicellular ancestors of animals; iii) *Monosiga brevicolis* represents choanoflagellates, unicellular and colonial organisms, considered the closest living relatives of animals. Among fungi, we did not find potential Ki-67 homologues in the model organisms of budding and fission yeast, but we selected other representatives: iv) *Malassezia globosa*, a yeast-like fungus, that causes dandruff and dermatitis; v) *Mycoemilia scoparia*, belonging to the order Kickxellales; vi) *Paramicrosporidium saccamoebae*, representing the Cryptomycota (‘hidden fungi’) clade of microorganisms that differ from classical fungi by the lack of chitinous cell wall. We included several organisms from outside the Opisthokonts clade: i) *Thecamonas trahens*, representing protozoan zooflagellates; ii) *Dictyostelium discoideum*, a soil-dwelling amoeba commonly known as slime mold; iii) *Hondaea fermentalgiana*, a marine unicellular saprotrophic eukaryote of the order Thraustochytrida, in the phylum Bigyra, which produces zoospores but is unrelated to fungi. The *Hondaea* protein might be incomplete as an Ensembl search reveals a long transcript annotated as FHA protein, and several short in-frame transcripts downstream. Entry of full-length human Ki-67 in SHOOT did not return any results in plants, but querying with human Ki-67 FHA domain alone did reveal plant proteins. The most closely related (45% identity with 5% of the sequence) was an FHA domain protein, described as a DNA helicase, of the single-celled green algae *Ostreococcus lucimarinus* (27415370_92541).

BLAST of the latter versus human taxid identified chain A of the PHD finger protein 21A (2PUY_A, 60 aa); the second hit was chromodomain helicase DNA-binding protein 4 (KAI2563986, 1905 aa), showing 39% identity with 7% of sequence. SHOOT also listed an *Arabidopsis thaliana* SMAD/FHA domain-containing protein (Q9M8A0, 585 aa), described as nuclear inhibitor of protein phosphatase-1. Neither show FHA domain residues characteristic of Ki-67. We conclude that there is likely no Ki-67 homologue in plants.

The model genetic organisms *Caenorhabditis elegans* and *Drosophila melanogaster* were also absent. However, SHOOT, which has superior sensitivity to local alignment ^13^ revealed a *C. elegans* protein (Q9N3H4; Supp. Fig. 2A) annotated as an FHA domain-containing protein which is a product of the Y53G8AL.1 gene. The WormBase database (https://wormbase.org/#012-34-5) describes the latter as encoding a hypothetical FHA domain-containing protein, whose mutation has no known phenotype.

Regarding fruit flies, the InterPro database contained a putative Ki-67-like protein from the fruit fly *D. willistoni*. To find a possible *D. melanogaster* protein, we used eggNOG (evolutionary gene genealogy Nonsupervised Orthologous Groups; ^20^), that employs both clustering and phylogenetic analyses to infer orthologous groups. Entry of human Ki-67 resulted in a list of 124 proteins, including non-vertebrates such as echinoderms, cnidarians, brachiopods, insects, arthropods, and others, and a *D. melanogaster* protein, under the PaxDB (protein abundance database) entry FBpp0288574. This corresponds to UniProt accession B7Z0I8, a 3441 aa protein, annotated as *D. melanogaster* Titin. This annotation is erroneous as BLAST with human Titin (a 34,350 aa long protein, involved in function of striated muscle and containing multiple Ig-like domains) against the *D. melanogaster* taxid returned a different 18,141 aa protein, with UniProt accession number Q9I7U4, consisting of multiple Ig-like domains, and thus clearly the Titin homologue. BLAST of B7Z0I8 against the *Drosophila* database (http://flybase.org/) confirmed the presence of a *D. melanogaster* protein encoded by Dmel/CG42232 gene. The protein has no known molecular function, belongs to the SMAD/FHA domain superfamily, and *Xenopus* and zebrafish Ki-67 are listed as its orthologues. Its peak expression is reported in the first 12 hours embryonic stages, indicating that it is initially maternally loaded and then also highly zygotically expressed (Supp. Fig. 2B), and during early pupal stages. We thus identified a putative *Drosophila melanogaster* Ki-67 homologue.

We compared the conservation of FHA domains of a subset of these putative Ki-67 homologues representing different phyla, with a set of FHA domains from human proteins unrelated to Ki-67 (see Fig. 2A). The phylogenetic tree reconstructed after MUSCLE sequence alignment showed clear segregation of FHA domains of the Ki-67-like proteins from FHA domains of unrelated proteins (Supp. Fig. 2C). This provided us with a baseline indication of relatedness of the identified candidate Ki-67 homologues.

### General features of Ki-67 proteins

Mammalian Ki-67 has several structural and sequence characteristics: its N-terminus is ordered and composed of the FHA domain; it is a large and predominantly intrinsically disordered protein; it contains a PP1-binding motif downstream of the FHA domain; it is highly basic at neutral pH; it has repeated sequences in its disordered part; and it contains many CDK phosphorylation sites (S/T-P) that become phosphorylated by CDK1 in mitosis, reversing the overall charge of the protein to negative at neutral pH. The majority of these features are present in our candidate Ki-67 homologues (Table 1). For taxa with potentially incomplete transcripts (e.g. *Hondaea*), the features are provisional, pending full-length validation. The location of the FHA domain is N-terminal in all homologues. Most have one predicted PP1-binding motif, [RK]-*X*_0 –1_-[VI]-{P}-[FW] ^21^, which is predominantly located in the N-terminus, although in some species it is found in C-terminus (e.g., *Phallusia*, *Drosophila*; Table 1); some species have two motifs. The size ranges from 632 aa for *Halobacteriovorax* to 4632 aa in mosquito, with an average length of 2000 aa. All are enriched in charged amino acids and most have a highly basic isoelectric point. AIUpred3 ^14^ predicts all proteins to be predominantly intrinsically disordered (51 to 89% overall disordered, with the exception of *Hondaea* at 40%, according to PONDR VLXT ^22^). CDK minimal consensus S/T-P sites exist in the disordered regions of all proteins, with frequency ranging from every 128 aa in *C. elegans*, up to every 8 aa in the scallop homologue (every 35 aa for human Ki-67), and in most proteins their phosphorylation would reverse the charge of the protein to negative at neutral pH. The RADAR repeat predictor ^16^ identifies repeated sequences in most of the analysed proteins. For vertebrate species, overall sequence homology with human Ki-67 ranges from 42% (blue whale) to 12% (shark), while for non-vertebrates, the homology is between 17% (starfish and lancelet) to 10% (*D. melanogaster*) (Suppl. Fig. 2D). Thus, even within vertebrates, the sequences are highly divergent, whereas in non-vertebrates, the homology is so low as to be unidentifiable were it not for the FHA domain. In summary, despite extremely limited overall homology, the identified proteins present similar structural and molecular characteristics to Ki-67.

### Analysis of cross-species conservation of the Ki-67 FHA domain

If the conservation of the FHA domain sequences of putative Ki-67 proteins from the different species reflects their common ancestry, then phylogeny of FHA domains should be similar to species phylogeny. To test this, we reconstructed a phylogenetic tree based on the sequence alignment of the FHA domains performed with MUSCLE (Suppl. Fig. 3A). As with the *Ciona* homologue, the alignment showed a striking overall conservation, with sequence homology with the human Ki-67 FHA domain ranging from 86% for blue whale to 17% for *C. elegans* (compare with the low sequence conservation between FHA domains of different human proteins; Fig. 2A). Next, a maximum likelihood phylogenetic tree was reconstructed using PhyML ^23^. The topology of the reconstructed tree was compared to that of the reference tree for the same set of species by generating a co-phylogeny that plots the NCBI and PhyML trees in mirror (Fig. 3A; see Methods for details). As expected, the results show that the orthology hypothesis clearly holds for the vertebrate clades in the NCBI tree, including human, whale, mouse, chicken, *Xenopus* and fish, and includes the cephalochordate fish-like invertebrate Lancelet, a close relative to vertebrates. Moreover, some distant clades (e.g. one including *C. elegans* and the arthropods silk moth, beetle, mosquito, *Drosophila*, barnacle, crab, *Daphnia* and tick) are also well conserved. We compared the NCBI and PhyML tree topologies and tested whether the distance between the two trees is compatible with what would be obtained by chance if the null hypothesis of no orthology were true by generating random trees. The topological distance was given by the maximum agreement subtree (MAST) criterion (Fig. 3B). This shows that random trees have, on average, about 15 tips in common with the reference tree. The FHA tree has 23 tips in common with the reference NCBI tree, thereby allowing us to reject the null hypothesis of no homology (p=3x10^-4^). It is therefore highly likely that these sequences are homologous.

**Figure 3.**
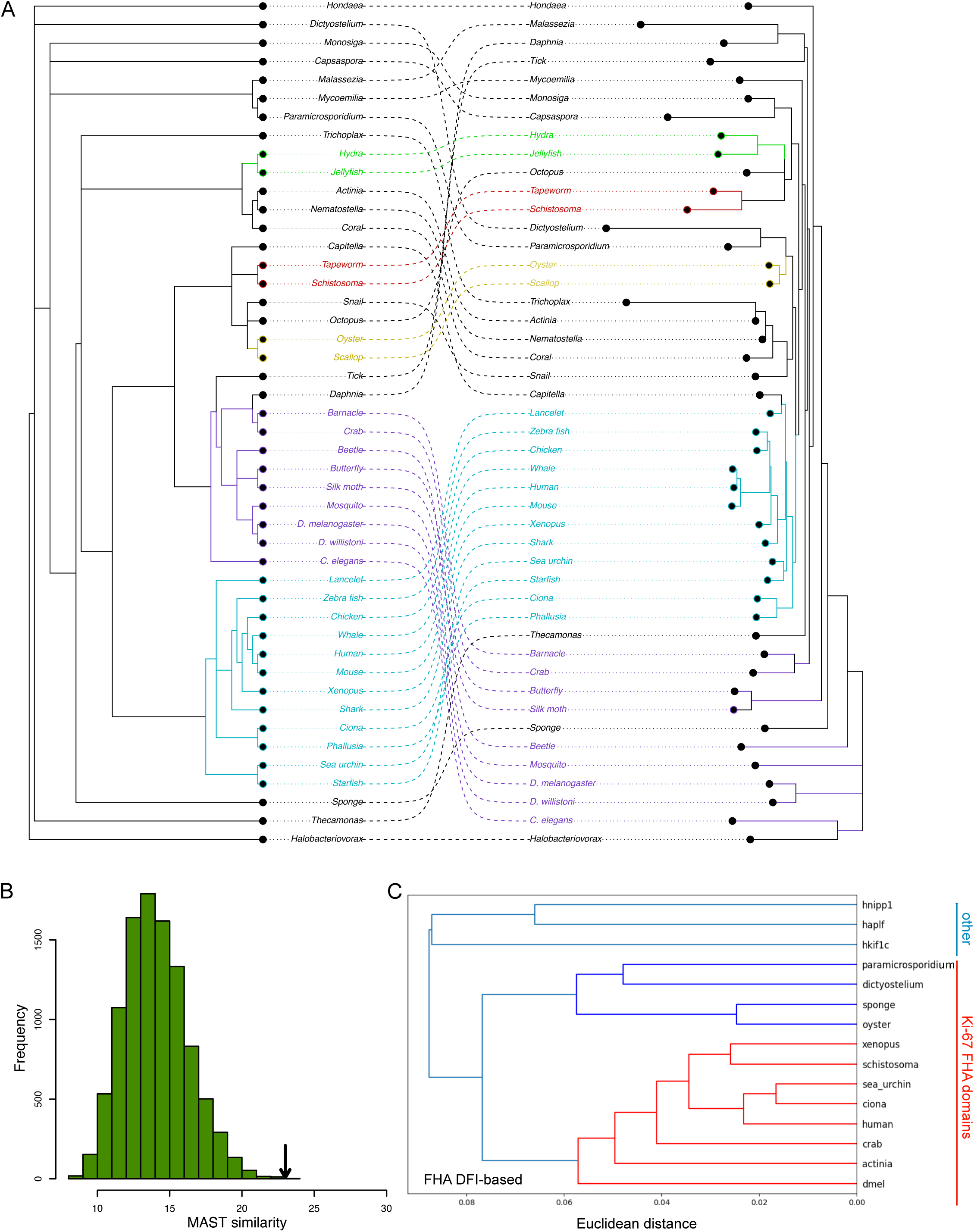
Sequence and dynamics of FHA domains allow assessement of homology. A. A co-phylogeny plotting in mirror the reference NCBI tree of species (left) and a maximum likelihood phylogenetic tree calculated using PhyML, based on MUSCL alignment of FHA domain sequences (right; here, the edge lengths represent expected number of substitutions between nucleotides, ie., the quantity of evolution). B. Comparison of NCBI and PhyML tree topologies expressed as MAST (maximum agreement subtree) criterion (see main text and methods); the arrow indicates the similarity between the NCBI and PhyML trees; p-value = 0.0003. C. Dendrogram of PCA-based clustering of raw Dynamic Flexibility Index profiles of a subset of FHA domains of Ki-67 homologues and unrelated human proteins (NIPP1, APLF, KIF1C). Seven PCs (accounting for 93.3% of the variance) were used.

To see if this conclusion holds true using a sequence-independent method, we assessed the relatedness of Ki-67 FHA domains and unrelated FHA domains by structural modeling. First, we used the root-mean-square deviation (RMSD) obtained from initial structure predictions generated by AlphaFold3. This showed large differences between FHA domains of putative Ki-67 homologues (especially *Paramicrosporidium*, sponge, and oyster) and also with the unrelated FHA domains (Supp. Fig. 3B). Thus, simple comparison of the initial structures does not unambigously distinguish Ki-67 and unrelated FHA domains. This is not surprising given that overall structure of FHA domains is highly similar, despite sequence divergence (see Fig. 2 A and B). We next evaluated the dynamic properties of the FHA domains of Ki-67 homologues by whole atom molecular dynamics (MD) simulations. We calculated the dynamic flexibility index (DFI; ^24,25^), that quantifies relative flexibility at single amino acid level; differences in DFI profiles have functional consequences (e.g. ^26,27^). A dendogram of PCA-based clustering of DFI profiles based on 2.5 µs of whole atom MD simulation (see Supp. Fig. 3C for raw profiles) shows that non-Ki-67 FHA domains cluster separately from the putative Ki-67 FHA domains (Fig. 3C). Interestingly, FHA domains from putative Ki-67 homologues clustered in two different branches, with the more divergent *Dictyostelium*, sponge, oyster and *Paramicrosporidium* belonging to a separate cluster from the human, *Xenopus*, *Ciona*, sea urchin, crab, anemone, *Schistosoma* and *Drosophila*, in line with the phylogeny that also showed higher divergence of these sequences (Fig. 3A).

We next explored whether the distinctive binding mode of human Ki-67 FHA domain with NIFK, its only known interactor, is conserved. AlphaFold3 modeling of these interactions across species showed that they fall into one of four behaviours (see Table 1; Suppl. Fig. 4A): i) The FHA domain interaction with the same species NIFK C-terminus, phosphorylated on existing TP sites, is conserved, including formation of the antiparallel β-strand in the NIFK and interaction of FHA domain loops with phosphorylated Thr (e.g. starfish, oyster, Monosiga). In the case of starfish, NIFK forms two β-strands interacting with the FHA domain. ii) The binding is similar to that described for *Ciona* (see Fig. 2E), with the formation of an anti-parallel β-strand in NIFK and phosphosites (or a negatively charged amino acid) interacting C-terminally of it (e.g. lancelet, *Phallusia*, *Schistosoma*). iii) FHA domain does not interact with NIFK of the same species, but interacts with human NIFK as does human Ki-67 FHA, indicating a conserved overall structure and function. In these species (e.g. butterfly, crab, *Daphnia*, *Hondaea*), NIFK often lacks the C-terminus and phospho-sites that mediate the interaction with Ki-67 FHA domain in human. In some species (e.g. tick, beetle, silk moth), the antiparallel β-strand forms with the human NIFK (conserved aa), but there is no interaction with ph-Thr. iv) For a few distant species, including *Drosophila*, *Capsaspora*, *C. elegans* and *Paramicrosporidium*, there was no specific binding with either the NIFK homologue or human NIFK, suggesting that the interaction of Ki-67 FHA domain with NIFK might have evolved after Ki-67 emerged or been lost in these species. No formation of the antiparallel β-strands between a phosphorylated peptide of NIFK and unrelated human FHA domains (CEP170, MDC1, KIF1C and CHK2) was predicted by AlphaFold3, although the interaction between the loops of these FHA domains and the ph-Thr can occur (with variable strength; Supp. Fig. 4B).

In summary, the FHA domain-level analysis confirms homology between the invertebrate putative Ki-67 proteins and human Ki-67. The remaining part of the protein is disordered, with no conservation of amino acid sequence. However, if the cellular functions of Ki-67 are conserved, the disordered regions might have conserved biophysical characteristics and/or sequence patterning.

**Figure 4.**
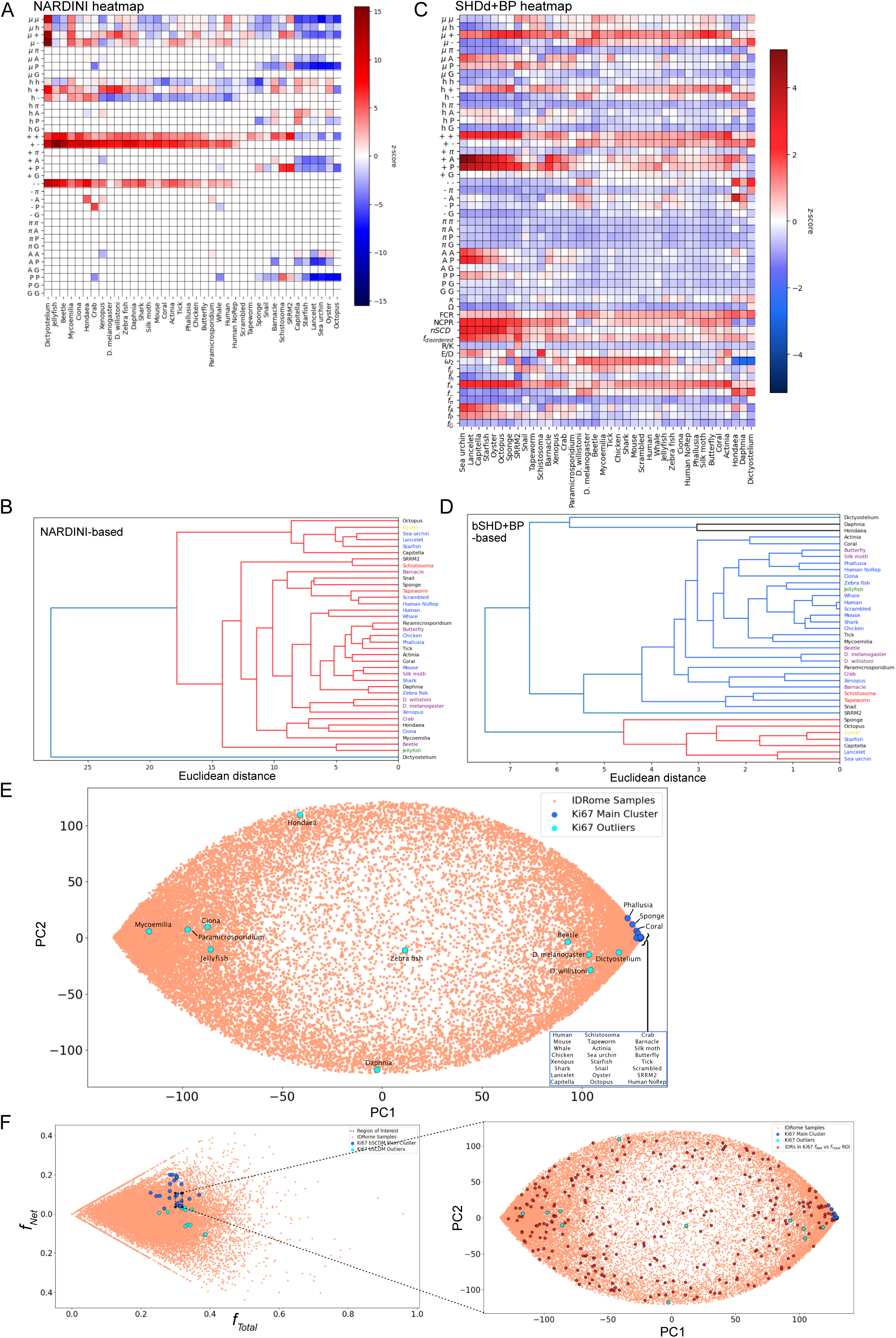
Sequence charge patterning and biophysical features show conservation between distant Ki-67 homologues and distinguish them from human IDRome. A. Heatmap of NARDINI features for Ki-67 sequences. Sorting is based on dendrogram clustering of NARDINI following PCA. Z-scores are derived from mean and standard deviation of 100,000 randomised sequences with the same amino acid composition but different patterning. Z-scores for blocky or well-mixed sequence features are shown as a red-to-blue colour scale. B. Dendrogram of NARDINI features following PCA (see methods) for Ki-67 sequences. Three principal components were required to account for 91.22% of variance. Species names are colored based on NCBI and FHA domain phylogeny (Fig. 3A). C. Heatmap of SHDd+BP features of Ki-67 sequences. The z-score value is averaged over the entire human IDRome. The sorting of the sequences is dictated by the dendrogram clustering of the SHDd distance matrix following PCA (D). D. Dendogram of SHDd+BP features following PCA (see methods). Five principal components were required to account for 90.67% of variance. E. PC2 vs PC1 plot of Ki-67 proteins and the full human IDRome using bSCDM as an input for PCA. Here, PC1 and PC2 account for 46.12% and 13.63% of variance in bSCDM, respectively. Ki-67 and IDRome bSCDMs were downsised to 200x200 to increase computational tractability. Salmon, IDRome proteins; blue, Ki-67 proteins being part of the central cluster; light blue, Ki-67 proteins seen as outliers. F. Left, fraction net charge (ƒ_net_) vs fraction total charge (ƒ_total_) for all Ki-67 proteins and IDRome proteins, with a region of interest (ROI) selected to enclose the maximum number of Ki-67 main cluster proteins. Right, bSCDM PCA-space plot, as in E, with human IDRs from the ROI identified and colored red. There are 274 IDRs and 11 Ki-67 main cluster proteins present within the ƒ_net_ vs ƒ_total_ ROI (left panel).

*The disordered regions of Ki-67 homologues share compositional and biophysical features.* IDPs and IDRs are characterised by heterogeneity of conformational ensembles that are encoded in the IDP sequence. Yet IDPs show low overall conservation of amino acid sequence, thus precluding assignment of function based on alignments. However, an ‘evolutionary signature’ or ‘molecular grammar’ of IDPs ^28,29^ proposes that certain molecular features and patterning are functionally relevant and conserved, and thus can be used to characterise IDPs and assess their functional homology. We therefore characterised the IDRs of Ki-67 homologues, using physics-based and computational tools, to determine whether they share such a common evolutionary signature.

To assess patterning features of the Ki-67 sequences, we employed the NARDINI (**N**on-random **A**rrangement of **R**esidues in **D**isordered Regions **I**nferred using **N**umerical **I**ntermixing) computational tool ^30^. This tool searches for IDR sequence patterns, such as linear clustering (positive values) or mixing (negative values) of 8 different classes of amino acids relative to one another, and compares it to 100,000 randomly scrambled sequences of the same composition, thus normalising for compositional bias. As internal controls we included a randomly scrambled sequence of human Ki-67, as well as human Ki-67 with a deletion of the repeat domain. Patterning of these two proteins can be distinguished from that of full-length IDR of human Ki-67, indicating that much of it is due to the repeats. The most striking feature of the NARDINI heatmap (Fig. 4A) relates to electrostatics: the clustering (denoted by the red colour) of charged residues (+;+), (+;-) and (-.-). This indicates that the clustering of basic and acidic residues is evolutionarily conserved and is likely of functional importance. A separate group of sequences (octopus, oyster, sea urchin, lancelet, starfish and capitella) show a non-random segregation (blue colour) of polar-polar (µ,µ), polar-hydrophobic (µ,h), (µ,+), (µ,P), and (P,P). We also included in the analysis the splicing factor SRRM2 (UniProt Q9UQ35), which does not have an FHA domain, but is similarly long, disordered, and basic (pI of 12.4). Like human Ki-67, SRRM2 can phase separate *in vitro* and *in vivo* ^31^, and is a CDK substrate whose phosphorylation is predicted to increase its propensity to phase separate ^32^. NARDINI analysis shows that SRRM2 shares similarities with Ki-67 but also features not observed in any Ki-67, such as clustering of polar residues with prolines, or clustering of prolines (observed only in *Schistosoma*), suggesting that these might be required for its function in nuclear speckles. A dendrogram of NARDINI features (Fig. 4B) showed that human Ki-67 clusters most closely with whale, similarly to the FHA domain (Fig. 3A). Vertebrate species often cluster with distant organisms (for example, mouse and shark cluster with silk moth, and *Xenopus* with both *Drosophila*), suggesting that convergent sequence patterning across distant taxa may enable analogous functional properties of the Ki-67 IDPs. In summary, NARDINI analysis demonstrates that charged clustering is a common feature across the majority of the Ki-67 homologue sequences and implies functional importance, as proposed for human Ki-67^33^.

To derive general biophysical properties that emerge from sequence patterning, we developed a physics-derived profiling of IDRs which we called **S**equence **H**ydropathy **D**ecoration **d**ecomposition (SHDd). SHDd relates to sequence-dependent interaction dictating conformation. This set of features includes interactions between all groups of amino acids (polar, acidic, basic, hydrophobic, aromatic, as well as P, A, G) and is directly computed from the sequence. This is compared against the entire human IDRome ^34^ to give a z-score for each parameter ^29,30^, with positive values (red) indicating more frequent occurrence, and negative values (blue) indicating under-representation relative to the IDRome mean and standard deviation of SHDd categories. A key difference from NARDINI is that SHDd scores depend not only on patterning (clustering or demixing), but also composition (higher or lower occurrence of a given amino acid type). These maps were successfully used to classify modern and ancestral sequences of NCBD, CID and their complexation ^35^. The second set encompasses 17 Bio-Physical (BP) features that are important for IDR conformation and function ^11,36–40^. These include kappa and omega parameters, which describe the segregation or mixing of, respectively, oppositely charged residues relative to each other, and charged residues and prolines relative to all other residues (higher values indicate clustering). Other parameters include: fraction of charged residues (FCR), net charge per residue (NCPR), normalised sequence charge decomposition (nSCD, that describes charge patterning), fraction of disordered residues, ratio of acidic residues (E/D), ratio of basic residues (R/K), sequence-dependent non-electrostatic interaction measure ω_2_, and fractions of different types of amino acids. Fig. 4C presents the heatmap of z-scores for the combined SHDd+BP profiling. Overall, SHDd profiling demonstrated a highly similar global profile of the disordered domains of Ki-67 across species. Several common interaction patterns become apparent. Pairing of polar and basic (µ,+) and basic (+,+) residues is conserved across all species, and to a somewhat lesser extent (h,+), (+,P) and (+,A). On the other hand, a clear underrepresentation of interaction between polar and hydrophobic with aromatic residues is observed, as well as acidic and hydrophobic with glycine. Vertebrate Ki-67 homologues do not display any strong differences compared with other organisms, and the clustering indicates patterning similarities with such distant species as *Drosophila*, silk moth or *Daphnia*. The SDHd pipeline thus underscored the importance of basic residues and patterning in Ki-67 homologues.

The BP profiling shows that most sequences display lower kappa and omega values relative to the human IDRome baseline (indicating a higher mixing of charged residues, and charged and proline residues relative to all other), with only a few exceptions. This translates into more extended chain conformations (which was confirmed by size scaling analysis, see below). Most sequences also show higher values of FCR, NCPR, fraction of disordered residues, and fraction of basic residues (due to enrichment in lysines, as R/ K ratio is low in all species). Aromatic residues and glycines are uniformly underrepresented. The non-electrostatic interaction measure ω_2_ is elevated in vertebrates, but also species as distant as *Drosophila*, but is lower in *Daphnia*, *Hondaea* and *Dictyostelium*. Finally, although patterning affects the profile, amino acid composition accounts for the majority of the profiling, as shown by the only marginal differences between human Ki-67 and the scrambled control.

In SHDd+BP-derived dendogram (Fig. 4D), highly related species were more closely clustered, yet, as with NARDINI, there was still considerable intermixing of clustering of Ki-67 homologues from distant species, suggesting much stronger conservation of amino acid composition and patterning, not visible in direct sequence alignment. Indeed, deletion of repeats from human Ki-67 clusters it differently, probably due to the lower fraction of acidic and increased fraction of polar residues, and reduction in hydrophobic interactions. Similarly to what we observed with NARDINI analysis, a separate cluster appears, grouping sponge, octopus, oyster, starfish, capitella, lancelet and sea urchin, characterised by high FCR, NCPR, fraction of basic residues, enrichment in prolines and alanines, and the highest values of nSCD, indicating well-dispersed positively and negatively charged residues, and thus larger dimensions (see also Fig. 5A). In the SHDd+BP profiling, SRRM2 appears at the margin of the cluster containing vertebrate Ki-67 proteins, presenting, similarly to NARDINI, a mixture of various features shared with different Ki-67 homologues. In contrast to Ki-67 homologues that all show decreased R/K ratio compared to the human IDRome, SRRM2 is the only protein with increased R/K ratio.

**Figure 5.**
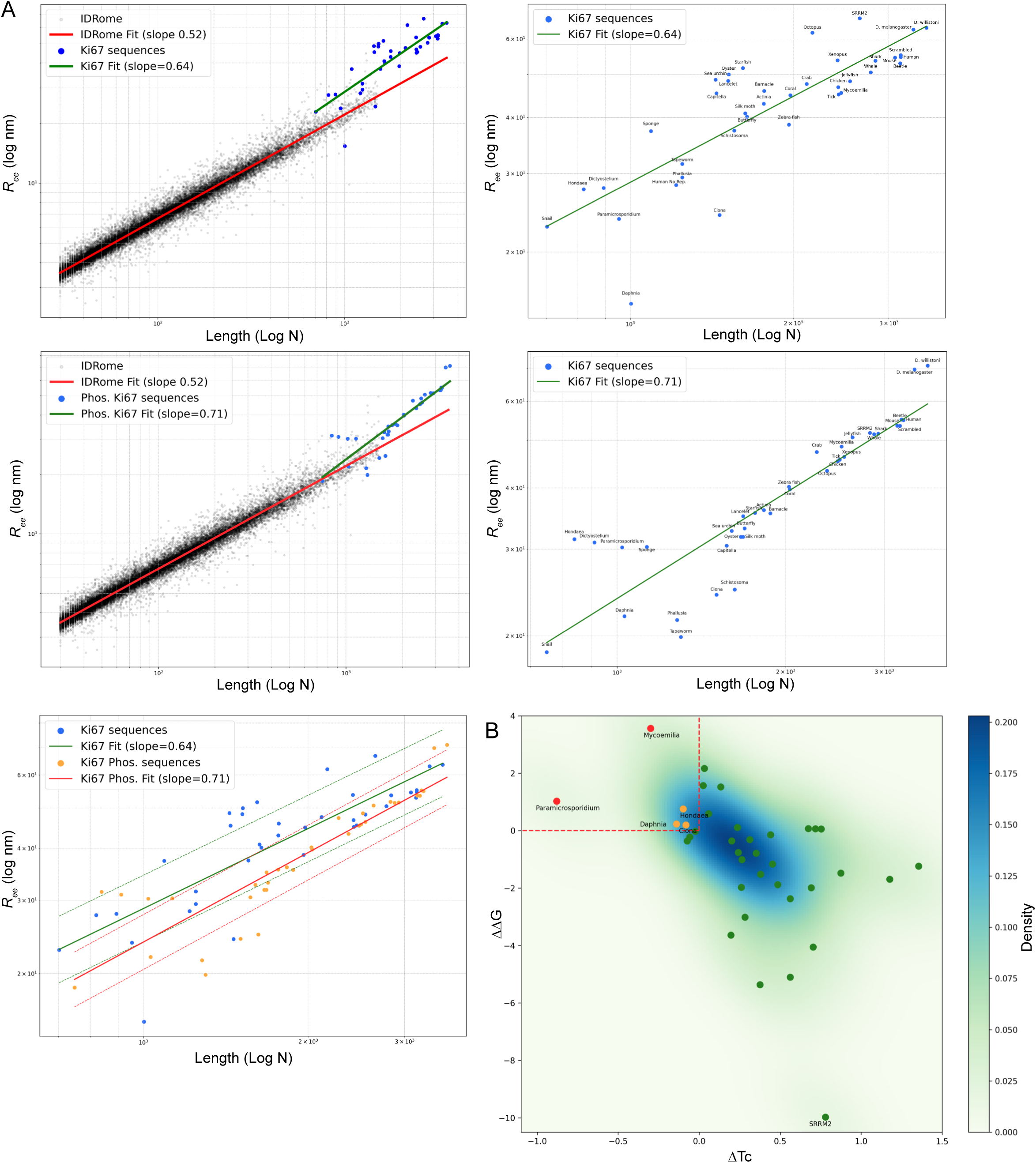
Ki-67 homologues are characterised by lower overall compaction and increase in phase separation propensity upon phosphorylation. A. Top left, scaling plot of the disordered proteome (un-phosphorylated; in black) with best fit line (red; slope of 0.52) and unphosphorylated Ki-67 homologues (in blue) with best fit line (green; slope of 0.64). Bottom left, scaling plot of the disordered proteome with phosphorylated Ki-67 proteins (blue; best fit line in green, slope of 0.71). Right, scaling plots of Ki-67 homologues with best fit line: top, unphosphorylated (slope of 0.64), and bottom, phosphorylated (slope of 0.71). Utmost bottom, scaling plot of Ki-67 non-phosphorylated proteins in blue with green best fit line and one sigma (0.080) dotted lines, and Ki-67 phosphorylated proteins in orange with red best fit line and one sigma (0.066) dotted lines. R_ee_, end-to-end distance. B. ΔΔG (phosphorylated ΔG - non-phosphorylated ΔG) vs ΔTc (phosphorylated Tc - non-phosphorylated Tc) Ki-67 plot, with outliers highlighted in orange (ΔΔG > 0 and ΔTc < 0) and extreme outliers highlighted in red (ΔΔG > 0.5 and ΔTc < −0.1).

Given that the most striking freature that emerged from NARDINI and SHDd+BP profiling was clustering and overrepresentation of charged residues, as well as high NCPR values, we next characterised Ki-67 IDRs using sequence charge decoration matrices (SCDM) _36,37_. SCDM was derived using a physics-based approach to describe sequence-specific interactions on chain conformation. These sequence-dependent interaction metrics dictates the distance between any two amino acid residues, and is predictive of many chain physics such as phase separation propensity ^12,41^. These metrics cannot be derived from simulation nor from statistics. We used binarised SCDM (bSCDM), that simplifies comparison of conserved biophysical features of sequences. To see how bSCDMs of Ki-67 sequences compare with other IDRs, we lowered the dimension of this matrix by PCA and compared against the entire human IDRome ^34^ (Supp. Fig. 5A and B) along with Ki-67 homologues (Fig. 4E). This showed that the majority of Ki-67 sequences cluster at one extremity of the IDRome PCA space, suggesting atypical shared features relative to most human IDRs. The bSCDM maps of the majority of Ki-67 sequences are characterised by predominantly repulsive interactions (Supp. Fig. 5C, I; bottom left corner of matrices, representing unphosphorylated proteins); the few outliers present mixed repulsive and attractive interactions (Supp. Fig. 5C, ii). The SCDM profile of SRRM2 is also highly repulsive and thus it clustered together with Ki-67 proteins. We wondered whether the highly repulsive interactions of most Ki-67 sequences are due to their high net positive charge. We thus plotted the net charge and the total charge of Ki-67 against the human IDRome space and found that Ki-67 homologues are scattered within the IDRome charge composition space (Supp. Fig. 5D). Thus, the high dimensional patterning of the charge (captured by bSCDM), rather than the total charge, distinguishes Ki-67 homologues from other IDRs. This was further confirmed when we selected a small sample of IDRs similar to most Ki-67 sequences in net charge vs total charge and plotted them against the PC (of bSCDM) IDRome space. These unrelated IDRs scattered throughout the bSCDM space (Fig. 4F), indicating that despite similarity in net and total charge, their charge patterning is distinct. In summary, consistent clustering across bSCDM, NARDINI and SHDd+BP features supports conserved higher order patterning beyond net/total charge of Ki-67, that likely underlie its critical roles in chromatin organisation. Moroever, we describe a methodological roadmap for multilevel analysis of IDR function and conservation.

### Ki-67 homologues are among the most highly extended proteins in the entire IDRome

IDP conformation is functional as it is conserved in orthologous sequences; it is an essential determinant of phase separation (PS) and formation of membraneless organelles ^34,42^, while compact IDRs are enriched in proteins binding chromatin and nuclear bodies ^34^. Proteins, as polymers, can be described in terms of scaling, the relationship between the global protein dimensions (expressed either as radius of gyration (R_g_) or end-to-end distance (R_ee_)) and the chain length ^43,44^. The scaling often reveals underlying physics of interaction and can be intrinsic to protein function ^45^. We therefore analysed the size scaling of Ki-67 homologues, in their non-phosphorylated and predicted phosphorylated states, in comparison with the entire human IDRome. To do so, we used our physics-based analytical model that accounts for charge patterning, while the non-charged patterning parameters were extracted from coarse-grained simulations and subsequently trained with machine learning ^46^. We could thus predict the ensemble average end-to-end distance of an IDP, accounting for both electrostatic and non-electrostatic contributions. This revealed that Ki-67 homologues are distinct from the human IDRome (Fig. 5A), both by their overall length and larger than expected R_ee_ (confirming the lower kappa and omega values observed in BP profiling, Fig. 4F). Next, since we have shown that CDK-dependent phosphorylation modulates IDP phase separation propensity of human Ki-67 ^32^, we determined the effect of phosphorylation of all S/T-P sites. This showed that phosphorylation further diminishes the differences between the Ki-67 homologues (SD of 0.066 vs 0.08 for non-phosphorylated sequences), and results both in higher protein compaction (decreased R_ee_) as well as a broader range of R_ee_. Thus, the more extended conformation is a common feature of diverse Ki-67 proteins, and allows regulation by phosphorylation.

### Phosphorylation increases phase separation propensity of most Ki-67 homologues

PS propensity is a property of human Ki-67 that appears important for its cellular function both in interphase and mitosis ^32,33,47^. Given the above results showing potential phosphorylation-regulated compaction, we assessed the PS propensity of Ki-67 homologues in their unphosphorylated and phosphorylated versions. To do this, we used a recent machine-learning model estimating the free energy change (ΔG) upon PS ^42^. We thus calculated the ΔG and saturation concentration for the putative Ki-67 homologues (Supp. Table 2). The ΔG was negative for most sequences, with the lowest values for beetle (-7), *Mycoemilia* (-4), tick (-3), and zebra fish (-2); therefore, these proteins are predicted to undergo PS without partners (based only on homotypic interactions) at a given condition of temperature and ionic strength _59_. Tapeworm, *Dictyostelium* and *D. willistoni* sequences had positive values of ΔG indicating that these proteins would not undergo spontaneous PS alone. To evaluate the effect of phosphorylation, we calculated ΔG for fully phosphorylated (on all CDK minimal consensus S/T-P sites) sequences. We found that the ΔG decreased (higher propensity for phase separation) for all proteins with the exception of *Mycoemilia*, *Paramicrosporidium*, and *Hondaea*, and to a lesser extent for *Ciona*, *Daphnia* and zebrafish (Supp. Table 2). Three proteins (sea urchin, crab and *Daphnia*) had positive values of ΔG even when phosphorylated. For 12 sequences, the ΔG was below -2, a value estimated as a threshold for spontaneous phase separation in isolation. The ΔG value for phosphorylated human Ki-67 was -1.5, thus not reaching the theoretical phase separation threshold ^59^; however, in our previous work ^32^ we showed that Ki-67 phase separates in cells when fully phosphorylated (upon inhibition of CDK-counteracting phosphatase), suggesting that the model underestimates PS in cells, where heterotypic interactions and crowding are involved. These data predict that phosphorylation of most Ki-67 homologues should promote phase separation.

To verify this conclusion by an orthogonal approach, we took advantage of our recently developed mathematical theory, renormalised Gaussian random phase approximation (rG-RPA _48_). This combines traditional RPA theory with sequence-dependent single-chain theory using a renormalised Gaussian (rG) chain formulation. We thus applied the rG-RPA model to analyse the PS propensity of Ki-67 homologues, calculating the critical temperature (T_c_; Suppl. Table 2). Intersection of the two approaches is shown in Fig. 5B. We plotted ΔΔG and ΔT_c_ (change in ΔG and T_c_, respectively, upon phosphorylation). Negative ΔΔG indicates that phosphorylation increases the propensity to PS, while the opposite is true for T_c_. There was a clear correlation between the two approaches. For most proteins, phosphorylation is predicted to increase PS propensity (lower ΔG and higher T_c_). Five sequences (*Mycoemilia*, *Paramicrosporidium*, *Hondaea*, *Ciona*, and *Daphnia*) did not pass the threshold of ΔT_c_ < -0.1 and ΔΔG < -0.5. We propose that merging the prediction of ΔG and the T_c_ is the most reliable approach to estimate the propensity to PS of proteins based on their sequence alone. Our analysis reveals that the majority of Ki-67 homologues have low propensity to phase separate when unphosphorylated, while for most of them the PS propensity should increase upon multisite phosphorylation by CDKs, suggesting similar PS regulation during the cell cycle. However, as noted above, this is an estimate only and the protein PS propensity will be strongly influenced by the cellular environment where heterotypic interactions and crowding are important.

### Ciona and Drosophila Ki-67 homologues differentially organise chromatin

Next, we tested experimentally whether Ki-67 homologues of two model organisms, *Ciona robusta* and *Drosophila melanogaster*, have conserved cellular localisation and function. To keep the cellular background constant and levels similar, we expressed the different homologues from the same vector in the same mammalian cell lines. These heterologous conditions allow direct comparison with human and mouse Ki-67 benchmarks. We first cloned the cDNA of CiKi-67 into a GFP-expression vector and transiently expressed it in human HeLa cells, which have prominent perinucleolar heterochromatin, and mouse 4T1 cells where pronounced chromocenters can be seen. As with human Ki-67 (Fig. 6A), in both cell lines, GFP-CiKi-67 localised to perinucleolar heterochromatin, distinct puncta in nucleoplasm, and chromocenters in 4T1 cells in interphase, and to the perichromosomal layer in mitosis (Fig. 6B and Supp. Fig. 6A). Like human Ki-67, at high levels of overexpression, CiKi-67 induced heterochromatin coalescence, indicating a conserved function in regulating chromatin organisation ^1,32^. These results indicate that the *Ciona robusta* Ki-67 is a *bona fide* functional homologue of human Ki-67, and, when expressed in mammalian cells, localises to the perinucleolar heterochromatin and PCL, and induces chromatin coalescence.

**Figure 6.**
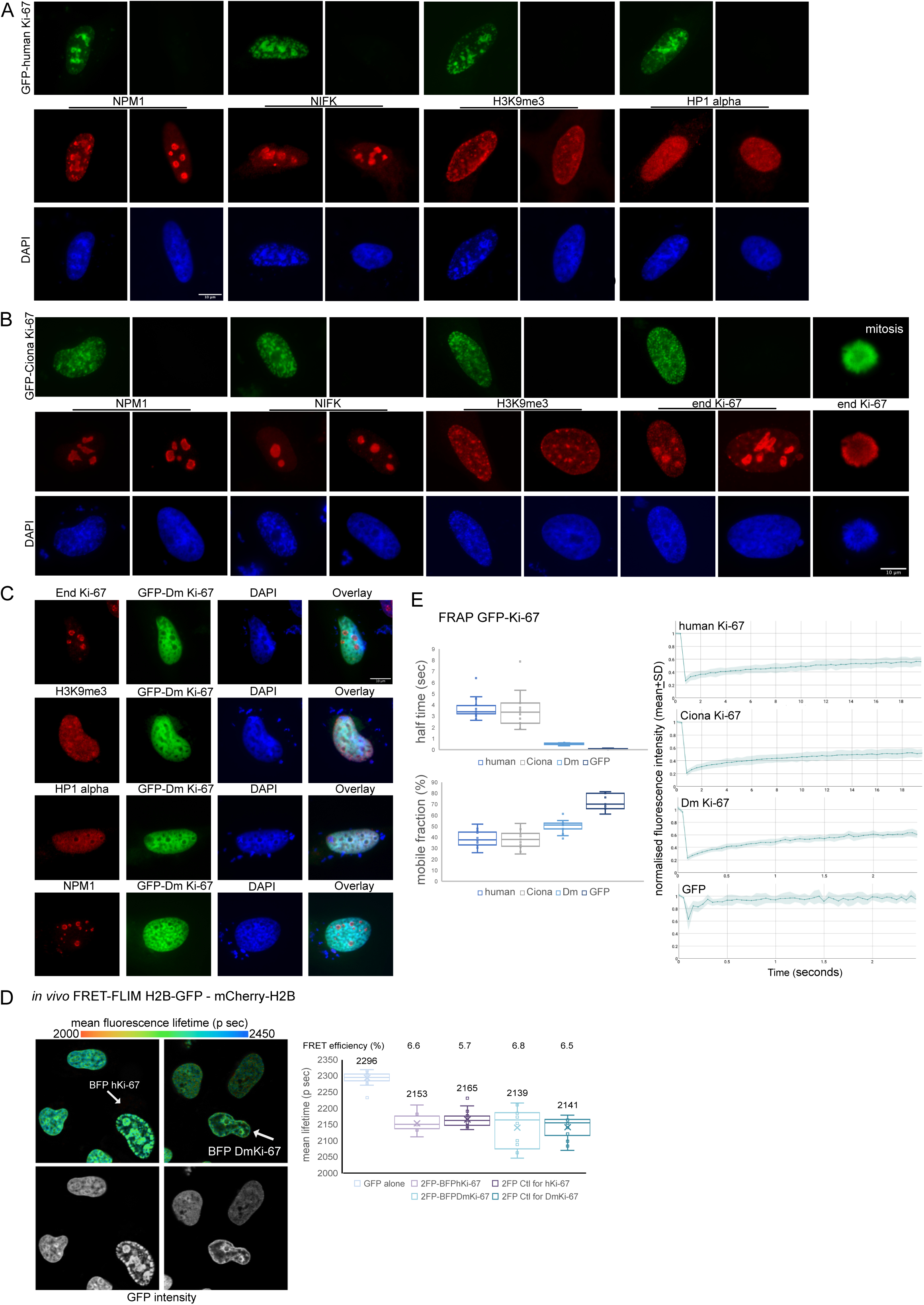
Ciona and *Drosophila melanogaster* Ki-67 homologues have conserved functions in regulating chromatin organisation. A, B, C. Immunofluorescence images of HeLa cells overexpressing GFP-tagged human (A), Ciona (B) and *Drosophila* (C) Ki-67, stained with the indicated antibodies; DNA was stained with DAPI. All cells are shown at the same magnification; scale bar 10 µm. D. FRET-FLIM analysis. Left, Box-and-Whisker plots representing mean fluorescence lifetime from HeLa cells expressing either the fluorophore donnor GFP-H2B alone (n=36; negative control for FRET) or H2B tagged with GFP and mCherry (2FP), overexpressing BFP-tagged human Ki-67 (n=12) or *Drosophila* Ki-67 (n=12); 2FP cells with no expression of hKi-67 (n=26) or with no DmKi-67 (n=22) from the same experiment were used as control. The horizontal lines represent the median, the middle crosses the mean, the boxes correspond to the fluorescence lifetime values from the 25-75^th^ percentile of the median, the whiskers cover the 10-90^th^ percentile values. The calculated FRET efficiency (%) is indicated above the graph. Right, representative images of live measurements of mean fluorescence lifetime (top) or GFP intensity only (bottom); cells expressing hKi-67 or DmKi-67 are indicated. Mean fluorescence lifetime is displayed using a continuous pseudo-colour scale from 2000 to 2450 p sec. E. FRAP analysis of the dynamics of human (n=20, 7 different cells), Ciona (n=15, 7 cells) and *Drosophila* (n=15, 5 cells) Ki-67. Left, half-time (sec) of recovery (top), and dynamic fraction (%, bottom). Right, mean normalised fluorescent intensity (+ SD) recovery curves. GFP was used for comparison (n=7, 3 cells). Representative of two independent experiments is shown.

Similarly to other Ki-67 homologues, the *Drosophila* Ki-67 (referred to as DmKi-67 thereafter) has an FHA domain in the N-terminal 100 aa, with the remainder being disordered, as predicted by AlphaFold3 and AIUpred (Supp. Fig. 6B). It also has repeated sequences within the IDR (Supp. Fig. 6B), and a PP1-binding motif (Table 1). However, it differs from vertebrate homologues by its highly negative charge (negative NCPR), although it presents similar clustering and patterning of charged residues (as shown by proximity to human Ki-67 in the IDRome bSCDM space, Fig. 4F). We assembled a full length 10kb cDNA from PCR-amplified fragments from *Drosophila melanogaster* larval cDNA, cloned it into the same GFP-expression vector as above and transiently expressed in HeLa cells. The overexpressed DmKi-67 induced reorganisation of the entire chromatin and, though it did not localise to nucleoli, caused relocalisation of the endogenous Ki-67 from its normal pattern (at perinucleolar heterochromatin and DAPI-dense foci in nucleoplasm) into nucleoli. This lead to nucleolar rounding up (Fig. 6C, Supp. Fig. 6C), suggesting a possible dominant-negative effect on endogenous Ki-67 that is in agreement with its displacement from heterochromatin. Consistent with this interpretation, the expression of the DmKi-67 in mouse NIH-3T3 cells led to strong reduction of DAPI-dense regions that correspond to chromocenters (Supp. Fig. 6D).

While overexpression of CiKi-67 recapitulates the coalescence of heterochromatin seen with human Ki-67, the nature of the reorganised chromatin induced by overexpressing DmKi-67 was less clear. Chromatin compaction can be assessed by the Förster Resonance Energy Transfer-Fluorescence Lifetime Imaging Microscopy (FRET-FLIM)-based assay in living cells stably expressing histone H2B tagged with GFP and mCherry ^49^. In this assay, interaction between GFP and mCherry of adjacent nucleosomes generates FRET, whose efficiency depends on the distance between the histones (FRET can take place only within < 10nm distance between donor and acceptor, and thus reflects nanoscale chromatin compaction). Using this assay, we have shown that ablation of Ki-67 in HeLa or mouse ES cells reduces heterochromatin compaction ^1,4^. To verify whether overexpression of Ki-67 alters chromatin compaction, we transiently transfected dual-tagged H2B HeLa cells with BFP-tagged human or DmKi-67. The results showed that overexpression of either Ki-67 did not alter average chromatin nanocompaction in living cells (Fig. 6D). Thus, the observed reorganisation of chromatin and coalescence of heterochromatin, that can be observed as DAPI-dense regions, does not result in increased proximity between nucleosomes. This is consistent with recent work in live cells where nucleosome proximity does not mirror the local DNA densisty ^4^.

A possible explanation for the differential effects on chromatin coalescence of human / *Ciona* Ki-67 compared to *Drosophila* might relate to their distinct isoelectric points. As DmKi-67 is negatively charged, whereas human and *Ciona* are positively charged, it might interact more weakly with chromatin, which could translate into higher mobility. Furthermore, human Ki-67 was shown to undergo phase separation in living cells ^32^, and one characteristics of phase separated proteins is their high dynamics as measured by FRAP (Fluorescence Recovery After Photobleaching). To characterise the protein mobility in cells, we performed FRAP in HeLa cells transiently expressing GFP-tagged human Ki-67, CiKi-67 and DmKi-67. This showed that human and *Ciona* proteins are highly and similarly dynamic, with t-half recovery of 3.7 and 3.5 sec, respectively, while DmKi-67 showed extreme dynamics (t-half of 0.5 sec; Fig. 6E). Given that protein diffusion coefficients are inversely proportional to their radii, yet the *Drosophila* protein has the longest chain length, these data indeed suggest that the interaction of the *Drosophila* homologue with chromatin is much weaker. The mobile fractions were similar (38%, 38%, and 49%), further indicating local interactions with chromatin and / or proteins. As expected, the control GFP protein was the most dynamic (t-half 0.08 sec, 72% mobile fraction). Overexpression of either human or DmKi-67 did not alter the dynamics of chromatin itself, as judged by FRAP of mCherry-tagged H2B (Supp. Fig. 6E). The histones remained predominantly immobile (mobile fraction of 9%, 3% and 5%, in control untransfected, GFP-human Ki-67, and GFP-DmKi67-overexpressing cells, respectively). Thus, diverse Ki-67 homologues show dynamics compatible with phase separation in living cells whereas chromatin itself does not, and IDR dynamics may be more influenced by charge than overall size.

## Discussion

Despite the accumulating knowledge of the important functions of Ki-67, the prevailing dogma has been that the protein is vertebrate-specific. This is enigmatic since the biological processes in which Ki-67 is thought to act are present throughout eukaryotes, while genome databases of non-vertebrates include sequences annotated as related to Ki-67. We identified Ki-67 homologues in all branches of animal kingdom. They are all large proteins that share a structured 100 amino acid-long FHA domain in their N-terminus, and which we distinguish from other FHA domains by structural modelling, phylogenetic and molecular dynamics tools. The proteins are predominantly disordered and we extracted a shared molecular signature of the Ki-67 IDRs. Finally, we cloned two homologues of Ki-67, which we subsequently overexpressed in human cells, and comparison of their phenotypes to human Ki-67 indicates both conserved and distinct functional properties.

The phylogenetic analysis of the FHA domains of Ki-67 homologues indicates common ancestry. Moreover, structured protein domains exist as dynamic entities, with atomic fluctuations, side chain rotations and whole domain movements, which is important for function ^25,50–52^. Consequently, analysis of the preservation of dynamic properties of residues critical for protein function can be used to study protein evolution ^24,53,54^. Indeed, analysis of the molecular dynamics of the FHA domain of Ki-67 homologues using dynamic flexibility index clustered them together and, importantly, distinguished them from unrelated FHA domains, confirming their relatedness.

The major part of Ki-67 is intrinsically disordered. Intrinsically disordered proteins, or proteins with intrinsically disordered regions constitute 20-40% of eukaryotic proteomes ^55,56^ and are increasingly recognised for their roles in almost all fundamental cellular processes, as well as in assembly and function of membraneless organelles ^57,58^. IDPs lack a fixed three-dimensional structure; rather, they exist as an ensemble of conformational states. They are also characterised by higher rates of sequence evolution than ordered proteins ^59^. Thus, if not known motifs or structured domains exist, searching for homologues using standard local alignment algorithms may fail. This difficulty may be alleviated by phylogenetic tools that assess sequence relationships within previously determined phylogenetic trees of protein families, increasing the sensitivity of homologue searches ^13,60^. However, decoding the sequence-function relationship of disordered domains remains challenging, underscoring a need for other approaches to determine evolutionary relationships between IDPs and to assign biological functions to them. Molecular dynamics simulations describe conformations of IDPs ^61–63^, but they do not necessarily allow comparison between non-homologous proteins and are computationally costly, prompting the development of AI-based models to predict conformation ^46,64–68^. Physics-based mathematical approaches that decode hidden features in sequences are also emerging. These account for sequence-specific electrostatic and non-electrostatic interactions and can provide functional information ^35–37,46,69,70^. Other strategies include analysis of compositional biases and statistics-based sequence patterns that may constitute an evolutionary signature, indicating functional relationships ^11,28–30,61,71–73^. In this study, we harnesed such existing tools and developed additional methods for analysing the patterning of the IDRs and comparing them to the entire human IDRome. We included biophysical parameters that constitute the so called ‘evolutionary grammar’ of IDRs and which relate to their function ^34,71–74^. We thus showed that despite the extremely low sequence homology, they share biophysical features that are conserved and thus likely important for their function. Ki-67 homologues present various conserved features, such as clustering of charged residues. The distinct charge patterning is captured by the highly dimensional metric of bSCDM (which has been previously successfully used for classifying functionally similar IDRs _36_), and which cannot be explained by simply analysing net and total charge of these IDRs. Most Ki-67 proteins cluster at one edge of the human IDRome bSCDM space, with few exceptions, which include the CiKi-67, despite the fact that the closely related *Phallusia* homologue is found in the main cluster, indicating some divergence between even closely related species. A noteworthy point is that oppositely charged proteins (e.g. the *Drosophila* and human Ki-67) show similar charge patterning, again indicating that the charge patterning is conserved independently of the nature of the net charge. Second, Ki-67 homologues share lower values of kappa and omega parameters, which indicates more extended chain conformations. This was confirmed by size scaling. Our strategy of combining multiple biophysical and patternign characteristics overcomes the difficulty of attributing functions from sequence conservation and provides a bluepring for assessing functional conservation between IDPs/IDRs.

In these analyses, we included the human SRRM2 nuclear speckle factor in the analysis, as it shares some characteristics with Ki-67 (is mostly disordered, is positively charged, can phase separate and is part of membraneless organelles), but does not contain an FHA domain. Interestingly, SRRM2 presents various features in common with Ki-67 IDRs. This reflects the remarkable co-evolution of SRMM2 with human Ki-67: it has a score of 3.16 (on a scale from 1-100) according to CladeOScope ^75^, indicating possible functional interactions. MUSCLE alignment revealed 21% identity between human Ki-67 and SRRM2 IDRs, which mostly concerned S, T, P and basic residues, suggesting that the amino acid composition is an important parameter of this co-evolution. Importantly, however, SRRM2 IDR also presents features that are not observed in any Ki-67 protein, such as high R/K ratio, while all putative Ki-67 homologues show low R/K ratio. This pattern likely underlies their distinct subcellular localisation, since K blocks are a feature of nucleolar proteins ^28^, while R patches are characteristic of nuclear speckle IDRs ^28,76^.

Finally, we cloned and studied in more detail two representants of invertebrate Ki-67, from *Ciona robusta* (belonging to tunicates, the sister group of vertebrates; ^77^) and *Drosophila melanogaster*. CiKi-67, when expressed in human or mouse cells, localises to the same nuclear regions as human Ki-67, is equally dynamic and can induce chromatin coalescence. The fruit fly Ki-67 homologue is also highly dynamic and reorganises chromatin. While chromatin structure is morphologically altered upon human, *Ciona* or *Drosophila* Ki-67 overexpression, its mobility remains low. Moreover, despite the reorganisation of chromatin by ectopic Ki-67, the chromatin nano-compaction does not change. This is in line with our previous study ^4^: proximity of nucleosomes is not proportional to visible DNA density. Therefore, coalescence of heterochromatin at high levels of human Ki-67 expression does not result in higher nano-compaction. Importantly, the presence of HP1 alpha, a known Ki-67 interactor, is required to maintain inter-nucleosome distances within heterochromatin ^4^. *Drosophila* Ki-67 additionally presents several different characteristics. It does not localise to nucleoli in human cells and may not interact with NIFK; rather, when expressed in human cells, it acts in some respects in a dominant-negative manner, displacing endogenous Ki-67 from heterochromatin and relocalising it into nucleoli. This behaviour is likely due to the oveall negative charge of *Drosophila* Ki-67 (pI of 4.5), which is expected to reduce interactions with RNA or with negatively charged nucleolar factors (*e.g.* NPM1), and likely also underlies its higher mobility. Nevertheless, the C-terminal domain, which is responsible for both DNA and HP1 interactions in human Ki-67 ^78,79^, is basic in both human and *Drosophila* (pI of 10.16 and 10.58 respectively), consistent with the conserved interaction with chromatin and regulation of its organisation. Divergences between vertebrate and invertebrate functions of IDRs have recently also been described for HP1 ^80^. While the yeast and *Drosophila* HP1 proteins phase separate and induce heterochromatin coalescence in mouse cells, the mouse homologue lost these properties, possibly because other proteins gained these functions. Ki-67 would be a good candidate to have gained at least some of the functions lost by HP1 during evolution, which might underlie the differences observed between *Drosophila* and human Ki-67. Alternatively, these might be linked to other genome regulation differences between invertebrates and vertebrates, for example the increasing presence and role in transcriptional regulation of DNA methylation (which is absent in yeast and *C. elegans*, and only trace levels are detected in *Drosophila*), which was accompanied by increased numbers of chromatin regulators binding methyl-CpG ^81^.

In summary, we identified Ki-67 homologues in a wide spectrum of non-vertebrate eukaryotes. Our results pave the way for studying Ki-67 homologues in tractable model systems, providing insights into conserved Ki-67 functions and mechanisms. Our strategy can be generalised to other IDRs that share a common domain, and our methodology provides an example of a roadmap to study their relatedness and conservation of functionally-essential features. Our multidisciplinary approach to comparative biology of intrinsically disordered proteins may therefore be useful for uncovering new functions of this significant fraction of the proteome.

## Materials and methods

### Sequence and structural analysis

Sequence alignments were generated using MUSCLE (MUltiple Sequence Comparison by Log-Expectation) multiple sequence alignement tool of EMBL and visualised with JalView (version 2.11.5.0; Waterhouse et al, 2009). Modeling of structures was performed using the AlphaFold3 web interface (Abramson et al, 2024). AlphaFold3 generates five predictions for each model, and the top ranked by AlphaFold3’s internal algorithms was selected for further analysis. Structural analysis and figure generation were performed using Open-Source PyMol (The PyMOL Molecular Graphics System, Schrödinger, LLC).

### Cloning

*Ciona* Ki-67-encoding sequence was amplified from VES80-F12 plasmid (kindly provided by J-P. Chambon, CRBM Montpellier) by PCR using Platinum SuperFi II DNA Polymerase (Invitrogen, #12361010), and cloned into pEGFP-C1 vector linearised by PCR by InFusion (In-Fusion® Snap Assembly Master Mix Takara, #638948) following manufacture’s instructions.

*Drosophila melanogaster* Ki-67 was cloned from cDNA of fly larvea kindly provided by J. Carnesecci (IGMM, Montpellier). In the first step, the sequence encoding the *D. melanogaster* Ki-67 was amplified by PCR in 10 fragments. Subsequently, the fragments were cloned into C1-EGFP vector in four steps, the three first ones (fragments #1 to #5) using InFusion (In Fusion® Snap Assembly Master Mix Takara, Cat.# 638948), the fourth (fragments #6 to #10) using HiFi cloning (NEBuilder® HiFi DNA Assembly Master Mix, 50 Reactions, Cat.# E2621L).

Human and *Drosophila melanogaster* Ki-67 were tagged with blue fluorescent protein (BFP) by replacing GFP in C1-EGFP vector with EBFP2 from Addgene plasmid 14893, by PCR.

### Antibodies

Ki-67- SP6, Abcam ab16667 (1:500)

NPM1 - SP236, Abcam ab183340 (1:1000)

NIFK - Invitrogen PA5-117374 (1:200)

MeCP2 - EPR23201-3, Abcam ab253197 (1:1000)

H3K9me3 - Abcam ab8898 (1:500) H3K9me2 – Abcam ab1220 (1:500)

HP1 alpha – clone 2HP-1H5-AS, Euromedex (1:500)

Goat anti-rabbit IgG (H+L) Cross-Adsorbed Secondary Antibody, Alexa Fluor™ 568, Invitrogen Ref: A-11011 (1:1000)

### Cell culture and transfections

All cells were cultured at 21% O_2_ and 5% CO_2_ in humidified conditions, in DMEM culture medium (Dulbecco’s Modified Eagle Medium, High glucose, GlutaMAX, 31966047, Life Technologies) + 10% fetal calf serum + 100 IU/ml-100 μg/ml penicillin-streptomycin. For transfections, cells were plated at 200 000 in 6-well plates 24h before. Transfection reagents used: JetPEI (Sartorius, # 101000053; DNA:JetPEI ratio 1:2); Lipofectamine^TM^ 3000 (Invitrogen L3000001; DNA:L3000 1:3); FuGene® HD (Promega, # E2311).

### Immunofluorescence

Cells were plated on coverslips (Decklglaser cover slips, Knittel Glass, d=10 mm). After 48 to 72 hours, depending on the treatment, cells were washed twice in PBS and fixed using 3.7% formaldehyde (Euromedex, 15714) in PBS for 15 minutes at room temperature. After a wash in PBS, cells were permeabilised with 0.25% TritonX-100 (Sigma Aldrich, T9284) in PBS for 5 minutes at room temperature. Cells were washed in PBS and blocked with 3% Bovine Serum Albumin (Sigma Aldrich, A3059) in TBS-0.1% Tween 20 (Sigma Aldrich, P7949) for 1 hour at room temperature. The coverslips were then incubated with primary antibodies resuspended in blocking solution for 1 h in a humidified chamber. After 5 washes in TBS-Tween, the coverslips were incubated with secondary Alexa Fluor® antibody at 1:1000 in blocking solution, for 1 h at room temperature. After 5 washes in TBS-Tween, the coverslips were incubated for 10 minutes with Hoechst 33342 (diluted in PBS), and after one more wash in PBS and one in MiliQ water, cover slips were mounted with ProLong Gold Antifade Mountant (Invitrogen, P36961), sealed with nail polish and imaged within one week.

### FRAP

HeLa cells were seeded on glass-bottom plates and transciently transfected with GFP-tagged human, Ciona and *Drosophila melanogaster* Ki-67, or with GFP only. Cells were imaged 24 h post-transfection, with the inverted confocal LSM 980 NLO microscope, operating on Zeiss Zen Blue software. Conditions of acquisition: 63x oil immersion objective, 4x zoom; 750V laser power, at 0.08%; time interval of 0.4 s for human and *Ciona* Ki-67, 0.05 s for Dm Ki-67 and GFP, 50 frames. FRAP was performed on a 1 µm^2^ region of interest, with two bleach iterations after 2 pre-bleach images. For FRAP of mCherry-H2B, 10 iterations of bleach were done, with time interval of 2 s, 50 frames in total. Analysis of FRAP data was done using the ImageJ FRAP plugin (https://imagej.net/plugins/frap-tools). Subsequent batch analysis was done with EasyFRAP web tool (https://easyfrap.vmnet.upatras.gr). Double normalisation method was used, followed by curve fitting using a single term exponential equation, to extract the T-half and mobile fraction from the curves.

### FRET-FLIM

FLIM-FRET experiments were carried out on a HeLa^H2B-2FP^ cell line which stably co-expresses histone H2B tagged at its C-terminus with GFP, and histone H2B tagged with mCherry at the N-terminus, as previously described ^49^. FLIM measurements were performed at 37°C with a 63x oil immersion lens, NA 1.4 Plan-Apochromat objective, with an inverted confocal LSM980 NLO microscope (Zeiss) equipped with an environmental black-walled chamber. GFP two-photon excitation was carried out at 890 nm by using a tuneable Chameleon Ultra II laser (tuning range from 680 to 1080 nm) that provided sub-150-fs pulses at an 80-MHz repetition rate. Detection of the emitted photons from excited GFP was achieved through the use of an HPM-100 module (Hamamatsu R10467-40 photomultiplier tube). Laser power was adjusted to give a mean photon count rate of about 2 × 10^5^ photons per second. The fluorescence lifetime imaging, corresponding to the time elapsed between laser pulses and the fluorescence photons detection, was provided by time-correlated single-photon counting (TCSPC) electronics (SPC-100; Becker & Hickl). Fluorescence lifetime measurements were acquired over 90s, and fluorescence lifetimes were calculated for each individual pixel in the field of view (512 × 512 pixels). FLIM analyses were performed using SPCImage software (Becker & Hickl). FRET causes a decrease in the fluorescence lifetime of the donor molecules (GFP). For FRET measurements, the mean fluorescence lifetime (τD) of the donor (H2B-GFP) was first determined under non-FRET conditions (in the absence of acceptor) by fitting the fluorescence lifetime decay with a mono-exponential model. In contrast, under FRET conditions, the experimental fluorescence decay curves were fitted using a bi-exponential model. The mean fluorescence lifetime of the donor in the presence of the acceptor (τDA; H2B-GFP with mCherry-H2B) was determined in 2FP-HeLa cells. To constrain the fit, the lifetime of the non-interacting donor population (τD) was fixed based on control measurements in HeLa cells expressing GFP-H2B alone. The conversion of fluorescence lifetime into FRET efficiency for each pixel in the images was achieved according to the formula: FRET efficiency = 1 − (τ_DA_/τ_D_).

### Orthology finding through phylogenetic coplots

We assume that the FHA domain of Ki-67 arose early during the course of evolution and was transmitted vertically so that orthologous copies are found in a broad range of organisms. According to this hypothesis, the phylogenetic tree reconstructed from these copies of the domain should closely resemble that of the reference species tree of the corresponding organisms. Protein sequences from 46 organisms ranging from bacteria to human were first aligned using MUSCLE ^82^. A maximum likelihood phylogenetic tree was estimated under the LG substitution model ^83^ with the FreeRate mixture model of rates across sites ^84^, using PhyML_23_. The topology of the reconstructed tree was then compared to that of the reference tree for the same set of species. The reference tree was obtained from the NCBI data base through the ETE toolkit ^85^. The R package phangorn ^86^ was next used to generate a cophylogeny that plots the NCBI and PhyML trees in mirror. The ordering of the tips in both phylogeny is chosen so as to maximise the agreement between the two topologies.

### Statistical test of orthology

We next compared the NCBI and PhyML tree topologies and tested whether the distance between the two trees is compatible with what would be obtained by chance if the null hypothesis of no orthology were true. More specifically, the topological distance was given by the maximum agreement subtree (MAST) criterion which corresponds to the number of elements in the largest set of species for which the two tree topologies are identical. We next simulated 10000 random tree topologies under Kingman’s coalescent model ^87^ and randomly assigned labels to the tips of the trees hence generated using the names of the species included in the present analysis. This labelling of tips amounts to generating trees with a phylogenetic signal that does not reflect evolution. The MAST distance was then evaluated between the NCBI tree and each random coalescent tree. Note that the obtained results would be identical for trees simulated under the birth-death process as the distribution of tree topologies is the same under the coalescent and birth-death tree generating processes. The histogram of MAST distances obtained under the null hypothesis was then compared to the MAST distance between the NCBI and PhyML trees and the corresponding p-value was calculated.

### Ki-67 IDR sequence processing and curation

Several of the theoretical methods applied here are suitable for modeling of intrinsically disordered proteins and not folded proteins. Therefore, to minimise the impact of highly structured regions on the analysis of IDRs, we analysed the disorder of Ki-67 homologues and removed structured regions. All Ki-67 proteins underwent this processing, with the exception of the scrambled human Ki-67 sequence, which was used as a control to assess the influence of local patterning on sequence properties, and SRRM2, which served as a Ki-67-unrelated control. Each Ki-67 sequence contains a highly conserved N-terminal Forkhead-associated domain. Because this domain is characterised by a well-defined fold, it was removed from all sequences prior to further analysis. Following that, sequence disorder was assessed using the IUPred2A predictor ^14^ to identify and mitigate the influence of long, highly structured regions downstream. The primary goal of this processing was to preserve the maximum sequence length possible while also removing significant structured elements. Structural trimming was restricted to C-terminal regions and was only performed if the ordered segment accounted for more than 15% of the total residues in the unaltered sequence. With this criteria, C-terminal structured regions were trimmed from two sequences: snail Ki-67 (residues 802 to 1186), and *Hondaea* Ki-67 (residues 920 to 1139).

None of the internal (other than the C-terminal end) structured regions were removed as they were a small fraction of the total chain. In the *Drosophila willistoni* Ki-67 sequence, a structured region (residues 837 to 923) was preserved due to its small relative size (2.4%) compared to the full 3620-aa sequence. Similarly, a short-structured region and a putative PP1-binding domain between residues 1110 to 1184 (3.3%) were retained in the octopus Ki-67 sequence.

### Computing Sequence Charge Decoration Matrix

Sequence Charge Decoration Matrix (*SCDM_ij_*) quantifies electrostatic interaction that dictate ensemble average distance between any two amino acids, *i* and *j*, for all *i* such that *i* > *j* + 1. An *SCDM_ij_* > 0 indicates electrostatic repulsion between residues *i* and *j*, whereas an *SCDM_ij_* < 0 indicates electrostatic attraction. *SCDM_ij_* can be constructed directly from a protein’s amino acid sequence, following:

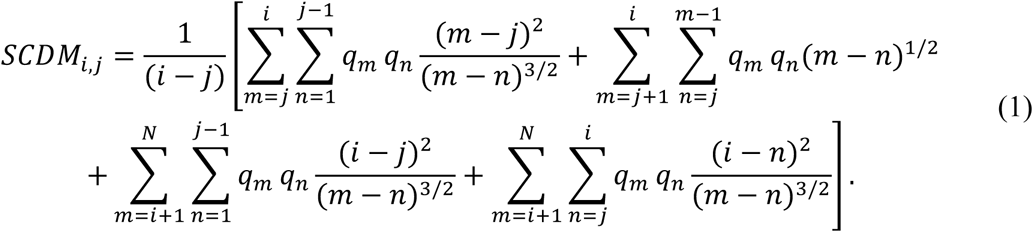

with, *q_i_* = −1 for amino acids *D*, *E*, *q_i_* = 1 for amino acids *R*, *K* and zero otherwise. Previous work has shown that the binarised form of *SCDM* can be used to classify functionally similar IDPs ^12,37,41^. Binary *SCDM* (termed *bSCDM*) helps capture the actual shape of the *SCDM* map which can be compared between proteins for functional classification. Binarisation is done by iterating through all *i*, *j* pairs of *SCDM* and assigning a new value based on the following rules: (1) if *SCDM* > 0, then *bSCDM*; is set to a value of 1. (2) if *SCDM* < 0 then *bSCDM* is set to a value of -1. (3) if *SCDM* is exactly zero, then *bSCDM* is also set to zero.

### bSCDM comparison between different proteins including the IDRome

Patterning information encoded in sequence charge decoration matrices (*SCDM*s) is intrinsically high-dimensional, and is difficult to compare across large number of proteins. We used principal component analysis (PCA) to reduce the dimensionality of these matrices and used principal components (PCs) across proteins including the IDRome (collection of all disordered proteins in the human proteome) for comparison. The overall scheme for the analysis across the IDRome was as follows. First, *SCDM*s were computed for all proteins in the dataset, including 27,060 IDRs comprising the IDRome and 38 Ki-67 protein sequences. We used a subsample of the entire IDRome ^34^ to exclude close homologues (see ^46^ for more). Because protein sequences vary substantially in length, the resulting *SCDM*s differed in dimensionality. To ensure uniform feature representations and maintain computational tractability, each *SCDM* was rescaled to a fixed size of 200 × 200. This resolution was empirically determined to be the largest vectorised matrix size for which PCA of the entire IDRome that could be computed reliably within available computational constraints. Matrix resizing was performed using the OpenCV Python package (https://pypi.org/project/opencv-python/), employing inner-area interpolation to preserve local patterning features while reducing dimensionality. This resizing kept the pattern of the actual map intact. Following resizing, each continuous-valued *SCDM* was converted into *bSCDM* as described above. Each 200 × 200 *bSCDM* was then flattened into a one-dimensional feature vector. Because *SCDM*s are symmetric by construction, only the lower triangular portion of each matrix was retained, excluding the diagonal, to avoid redundant information. This resulted in a fixed feature length of (200 × 199)/2 = 19,900 elements per protein. All flattened *bSCDM* vectors were assembled into a single global protein matrix of dimensions *n* × *p*, where *n* = 27,098 proteins (27,060 IDRs plus 38 Ki-67 proteins) and *p* = 19,900 features. PCA was performed on this global protein matrix using the scikit-learn Python package ^88^. The resulting principal components capture the dominant modes of variation in binary charge patterning across the combined IDRome and Ki-67 datasets. For visualisation, proteins were projected into principal component space, and the first two components were plotted as PC2 versus PC1. This two-dimensional representation enabled direct comparison of charge patterning of Ki-67 proteins in the background of the entire IDRome and facilitated identification of shared or distinct charge patterns within the PC space.

### Computing Sequence Hydropathy Decoration decomposition (SHDd)

*SCDM* as described above, provides a set of sequence dependent electrostatic patterning metrics arising from electrostatic interaction contributions to distance maps. A similar set of metrics arise from non-electrostatic (hydropathic) interactions and are denoted by the matrix called Sequence Hydropathy Decoration Matrix (*SHDM*). However, quantifying *SHDM* requires knowledge of the non-electrostatic interaction between all possible combinations of amino acids, which is difficult to measure or predict. In an earlier work (3), we showed that this matrix can be extracted by combining theory and all-atom simulation. However, for proteins considered here, all-atom or even coarse-grain simulation is not feasible due to their large size. To circumvent this, we use a different set of non-electrostatic patterning metrics called *SHDd*, also introduced in an earlier work that was shown to be useful in classifying proteins with identical charge patterning or low charge. *SHDd* is defined as below (see ^41^ for the details of derivation).

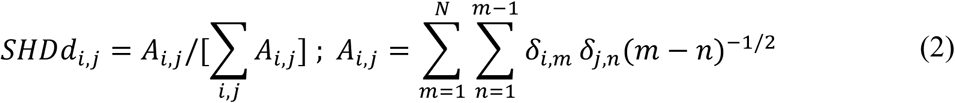

where, *i*, *j* are different types (or classes) of amino acids and *m*, *n* are amino acid indices along the protein backbone. The quantity *A_i,j_* represents a binary patterning metric for residue types *i* and *j*, which may be either distinct or identical. For simplicity we reduce the amino acid types/alphabet from twenty canonical amino acids to eight by grouping amino acids according to physicochemical similarity into the following categories: polar (S, T, N, Q, C, H), hydrophobic (I, L, M, V), acidic (E, D), basic (R, K), aromatic (F, W, Y), as well as an alanine, proline, and glycine group. This classification follows the scheme introduced by the Pappu group ^30^ and yields 8 × 8 dimensional *A* matrix and hence *SHDd* matrix, with 36 unique pairwise combinations. For comparative sequence analysis, it is advantageous to normalise these values against a reference distribution and compute a z-score, as done in previous studies^11, 30^. We have chosen the reference to be the human IDRome ^34^. The z-score is computed according to

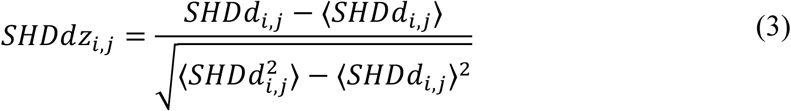

where ⟨. . ⟩ denotes averaging over the IDRome dataset. In summary, the *SHDdz* matrix encodes the binary patterning of each pair of amino acid groups (identical or distinct) relative to the background statistics of the human IDRome. This normalised matrix is used throughout the analyses presented in this work. We also combined *SHDd* with biophysical metrics (BP) for further analysis. BP, which was partially adapted from ^11^, consists of composition of the eight classes of amino acids and also the following metrics: kappa and omega, which are patterning metrics describing residue-residue interactions within chemically relevant amino acids groups ^89^, fraction of charged residues, net charge per residue, fraction of disordered residues, *ω*_)_ (a parameter describing non-charge interactions between residues; see section titled ‘Predicting dimensions of Ki-67 against the background proteome’ for more details), E to D ratio, and *nSCD* (see below for definition). Also included in BP is R to K ratio, which is useful since differences in LLPS propensity have been observed based on the identity of the basic residues (R or K) present in the sequence, not just from overall basic residue content ^90^.

*nSCD* is defined as,

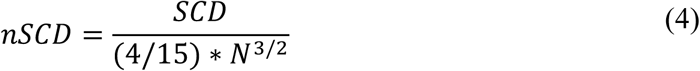

where *SCD* is sequence charge decoration ^70^, and *N* is the length of the protein. *nSCD* normalises *SCD* for chain length to allow comparison of charge patterning between proteins with different lengths.

### Computing non-random binary patterning metric using NARDINI

While *SHDd* – derived from statistical physics – captures patterning and maps sequence to interaction which is used to calculate conformation, an alternate (statistics based) set of patterning metrics is given by NARDINI (Non-random Arrangement of Residues in Disordered Regions Inferred using Numerical Intermixing). We applied NARDINI framework to analyse Ki-67 sequences. All calculations were performed using the localCIDER software package available from the Pappu Laboratory GitHub repository (https://github.com/Pappulab/localCIDER) ^30^. NARDINI generates a matrix of z-scores, in which each element quantifies the degree of non-random arrangement between two amino acid groups (which may be identical or distinct). For each element, the observed value is shifted by the mean and normalised by the standard deviation of a distribution derived from 100,000 randomised sequences of identical length and amino acid composition. Note, this z-score calculation is different from that of *SHDd* where the averaging is done over the entire IDRome. Amino acids were grouped into the same eight physicochemical categories as before: polar (S, T, N, Q, C, H), hydrophobic (I, L, M, V), acidic (E, D), basic (R, K), aromatic (F, W, Y), alanine (A), proline (P), and glycine (G). This grouping yields an 8 × 8 patterning matrix with 36 unique elements, including both diagonal and off-diagonal terms. For each Ki-67 sequence, the NARDINI analysis was conducted using the following default settings: (1) The z-score for each residue-pair category was computed using 100,000 randomised sequences for background distribution generation. (2) The randomisation seed was initialised using a floating-point value derived from the system date and time at the moment of computation. (3) The amino acid grouping was set to 8 × 8. The 36 unique elements of each z-score matrix were then extracted to produce a one-dimensional feature vector for every sequence.

### Computing Z-score of features

*SHDd* with BP and NARDINI all give high-dimensional features and some features may be redundant. Therefore, we perform PCA when using one dimensional feature vector extracted from *SHDdz* or NARDINI for consistency and dimensionality reduction. This process is primarily the same as described in the section above except resizing is not needed since *SHDdz* and NARDINI produce uniformly sized outputs regardless of protein size. The number of principal components retained was chosen to explain at least 90% of the total variance in the dataset. Each protein was then represented by its projection vector in this reduced PC space, and pairwise Euclidean distances between these projected vectors were computed to quantify sequence relationships as described below.

### Constructing distance matrices and dendrograms

All dendrograms reported here were generated from square pairwise distance matrices of size *n* × *n*, where *n* corresponds to the total number of protein sequences analysed. Each matrix element (*d_i,j_*) represents the quantitative dissimilarity between protein sequences *i* and *j*. The definition of this dissimilarity metric depends on the specific feature vector employed. For all analysis types, PCA was first applied to the corresponding feature matrices. Pairwise distances were then computed as Euclidean distances between the principal component vectors associated with each protein sequence. These distances served as the basis for constructing the corresponding distance matrices (*d_i,j_*). Hierarchical clustering was performed on each distance matrix using the average method implemented in the SciPy hierarchical clustering package ^91^. The resulting distance matrices were used to construct dendrograms depicting the hierarchical relationships among protein sequences for a given set of features.

### MD simulation protocol

All-atom molecular dynamics (MD) simulations were initiated from models generated by the AlphaFold3 server ^18^. All simulations were performed with the Amber package using the ff19SB force-field and the OPC water model ^92–95^. Minimisation and equilibration were performed using a standard protocol that has been described in earlier work ^96^. For the FHA domains, a 2.5 *μ*s equilibration simulation was performed first on the entire protein sequence. Clustering was performed on the last 2 *μ*s of the trajectory using the hierarchical agglomerative clustering method. Based on the simulation, we found that disordered tails at the end of the FHA domain can influence the Dynamic Flexibility Index (DFI, see below for more) profile of the folded domains. Since the goal is to compare DFI profiles of the folded domains across sequences to seek their similarity, it is important to minimise these interactions. Thus, we further trimmed the original sequences to remove residues predicted to be disordered. For each sequence, a representative structure from the dominant cluster of the MD simulation was selected, and only the first 100 residues were retained; the remaining C-terminal residues were deleted. These trimmed structures served as starting points for a second round of MD simulations. As controls, we used FHA domains of human APLF, KIF1C, and NIPP1 (UniProt Q8IW19, aa 4-103; O43896, aa 498-597; Q12972, aa 28-127, respectively) which were selected to match the 100-aa length of Ki-67 FHA domains; the disordered residues were trimmed based on the AlphaFold3-predictions. For all proteins (trimmed FHA domains of Ki-67 homologues and controls), minimisation and equilibration were performed as before, followed by a 2.5 *μ*s production simulation. The first 0.5 *μ*s of all production simulations was further discarded as equilibration time. The covariance overlap was calculated as described in _96,97_ and ^98^ for the last 1 *μ*s and 2 *μ*s demonstrating sufficient self-consistency in the simulations. This trajectory was then used to compute the DFI profile as detailed below.

### Calculating DFI

The Dynamic Flexibility Index (DFI) measures how a given position responds to external perturbations at other distal locations in the protein, thereby quantifying how sensitive individual residues are to global structural dynamics. Positions with high DFI values respond more to these perturbations due to their flexibility, while positions with low DFI values are more rigid and instead transfer the effects of the perturbation to the rest of the protein. The method for obtaining DFI profiles has been described in depth in previous work ^26,27,99,100^. We utilised 50 ns intervals, with a 5 ns overlap for computing covariance matrices throughout the final 2.0 μs of the simulation. We also computed DFI profiles using 25 ns, 75 ns, and 100 ns sized windows (see Supp. Fig. 7) to demonstrate the robust nature of the DFI profiles. We again used PCA when DFI profile was used as a feature to classify sequences. The protocol for PCA, distance matrix and dendrogram generation were identical to the one described above for other features.

### Predicting dimensions of Ki-67 against the background proteome

We computed dimensions of Ki-67 sequences using previously derived sequence-dependent free energy function of an IDP as a function of the chain dimension variable *x* _37,101_ given by:

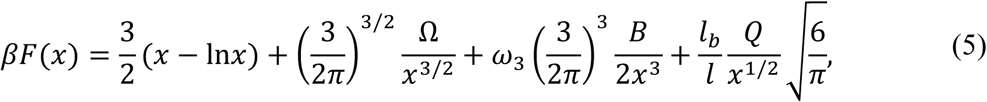

The first, third and the fourth terms can be computed directly from the sequence (using a known value of *ω*_3_: 0.2); see ^98^ for definition of Ω, B and Q. However, the second term – quantified by the sequence-dependent non-electrostatic interaction parameter *ω*_2_ cannot be directly obtained from the sequence and is defined as follows:

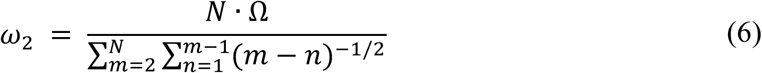

We developed Physics-Based Machine Learning (PML) ^46^ which employs a machine learning model to predict *ω*_2_. Using this previously trained model we predict *ω*_2_ for an unknown sequence. With the knowledge of *ω*_2_, we minimise the free energy to determine the most likely *x* for a given sequence. The ensemble average end-to-end distance 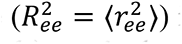 is calculated from *x* using 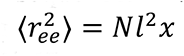 where, l=5.5Å (see ^37^ for more). Using this method, we computed 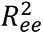 for the Ki-67 sequences and the IDRome and plot these values against *N* in log-log space. We calculated the best fit line and the 1*σ* standard deviation of the plotted points from the best fit line.

### Predicting LLPS propensity using machine learning

We used a machine learning based tool to predict phase separation propensity of disordered proteins directly from the sequence ^42^. We predict 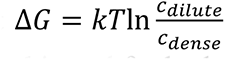 (*c_dilute_*, *c_dense_* are the concentrations of the dense and dilute phases) using this tool for both the non-phosphorylated and phosphorylated Ki-67 sequences. A high value of Δ*G* indicates the dilute and dense phases have significant differences in their density (*c*), implying a higher propensity to phase separate.

### Predicting critical temperature using a sequence-dependent physics-based theory of phase separation

While the above gives a measure of differences in the density between two phases, it does not give the critical temperature (*T*_0_) nor the phase diagram. Critical temperatures provide another measure of phase separation propensity, higher *T*_0_indicating a higher propensity to undergo LLPS. We use a polymer physics-based theoretical formalism to calculate sequence dependent *T*_0_. This formalism builds on our earlier model ^48^ of LLPS in IDP solutions accounting for the exact placement of charges in the sequence. We used Random Phase Approximation (RPA) and combined with sequence-dependent chain conformations (using equation), giving rise to a theory called rG-RPA (renormalized Gaussian RPA; see ^48^ for details) that accounts for spatial variations in density (density fluctuations) and sequence yielding solution free energy (*βf*) as,

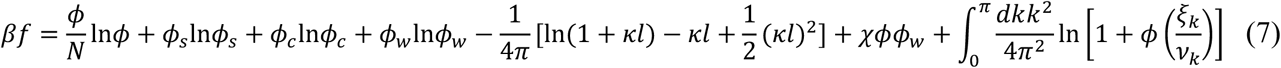

where the first four terms are the entropies of mixing with *ϕ*, *ϕ_s_*, *ϕ*_0_, and *ϕ_w_* = (1 − *ϕ_s_* − *ϕ_c_* − *ϕ*) being the volume fractions (related to density) of the polymer, salt, counterions, and water, respectively. The fifth term is the free energy of small ion interactions, determined by the debye screening length, 1/*κ* = 1/(4*πĩ*_b_.(*ϕ_s_* + *ϕ_c_*))^1/2^;*k* is non-dimensional wave vector; *ν_k_* = *k*^2^/(4*πĩ*_B_) + (*ϕ_s_* + *ϕ_c_*) is the inverse of the Fourier transform of screened coulomb potential accounting for salt effect; *ĩ_B_* (non-dimensional Bjerrum length) is inversely proportional to temperature; *N* is the number of amino acids. The sixth term is the usual mean-field interaction, the same as that in the Flory Huggins theory ^102^, with *χ* being an effective interaction term. The last term captures interaction that goes beyond mean-field Flory-Huggins (capturing density fluctuations) and embeds dependence of charge-sequence in *ξ_k_* and *ξ͞_k_*, defined as,

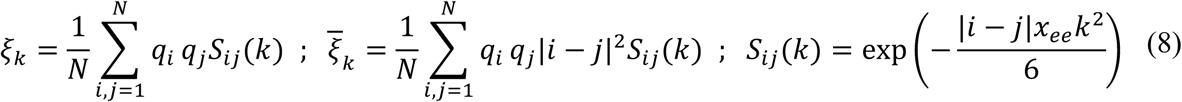

where, *q_i_*s are charges (1 for arginine, lysine, -1 for aspartic, glutamic acid, and 0 for others). Phosphorylation of serine or threonine is modeled by assigning *q* = −2 to mimic the effect of double negative charge of phosphate groups, and *q* = −1 for phosphomimic amino acids (aspartate/glutamate). *x_ee_* is sequence dependent and solved from equation,

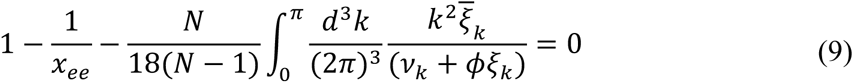

where *ξ_k_*, *ξ͞_k_* are defined above.

We rigorously tested rG-RPA against existing data on multiple charge systems (see ^48^ for more detail) as it explicitly models sequence charge dependence. However, so far, we assumed *χ* accounting for non-charge interaction to be sequence-independent. We now use these sequence-dependent *ω*_)_ values described above (equation [5]) to model *χ*. Specifically, we use *χ* = 0.5(1 − *ω*_2_ ∗ 310/*T* where, *T* is temperature. This relation is well known in polymer physics ^102^ when relating single chain behavior to that of many chains (to model solution). Thus, the theory above accounts for sequence-dependent charge interaction with fluctuations and sequence-dependent non-charge interaction at a mean-field level (without density fluctuation) from which phase diagrams along with critical points can be calculated.

## Acknowledgements

We acknowledge D. Coudreuse (IBGC, Bordeaux), M. Lagha (IGMM, Montpellier), F. Erdel (CBI, Toulouse), J. Dejardin (IGH, Montpellier), and R. Feil (IGMM, Montpellier) for valuable criticism of the paper; N. Lamb (University of Denver, US) for discussion and input in building LLPS model. We acknowledge the MRI imaging facility (BioCampus Montpellier), member of the national infrastructure France-BioImaging (https://ror.org/01y7vt929), supported by the French National Research Agency (ANR-24-INBS-0005 FBI BIOGEN). This work was supported by La Ligue contre le cancer (grant EL2025.LNCC/DaF: DF lab and AZ); the University of Montpellier LabMUSE EPIGenMed collaborative research project 2022 (GD); Fondation pour la Recherche Medicale (doctoral contract ECO202506050227 for AZ); NIH grant R01GM138901 (KG).

---

**Supplementary Figure 1.**
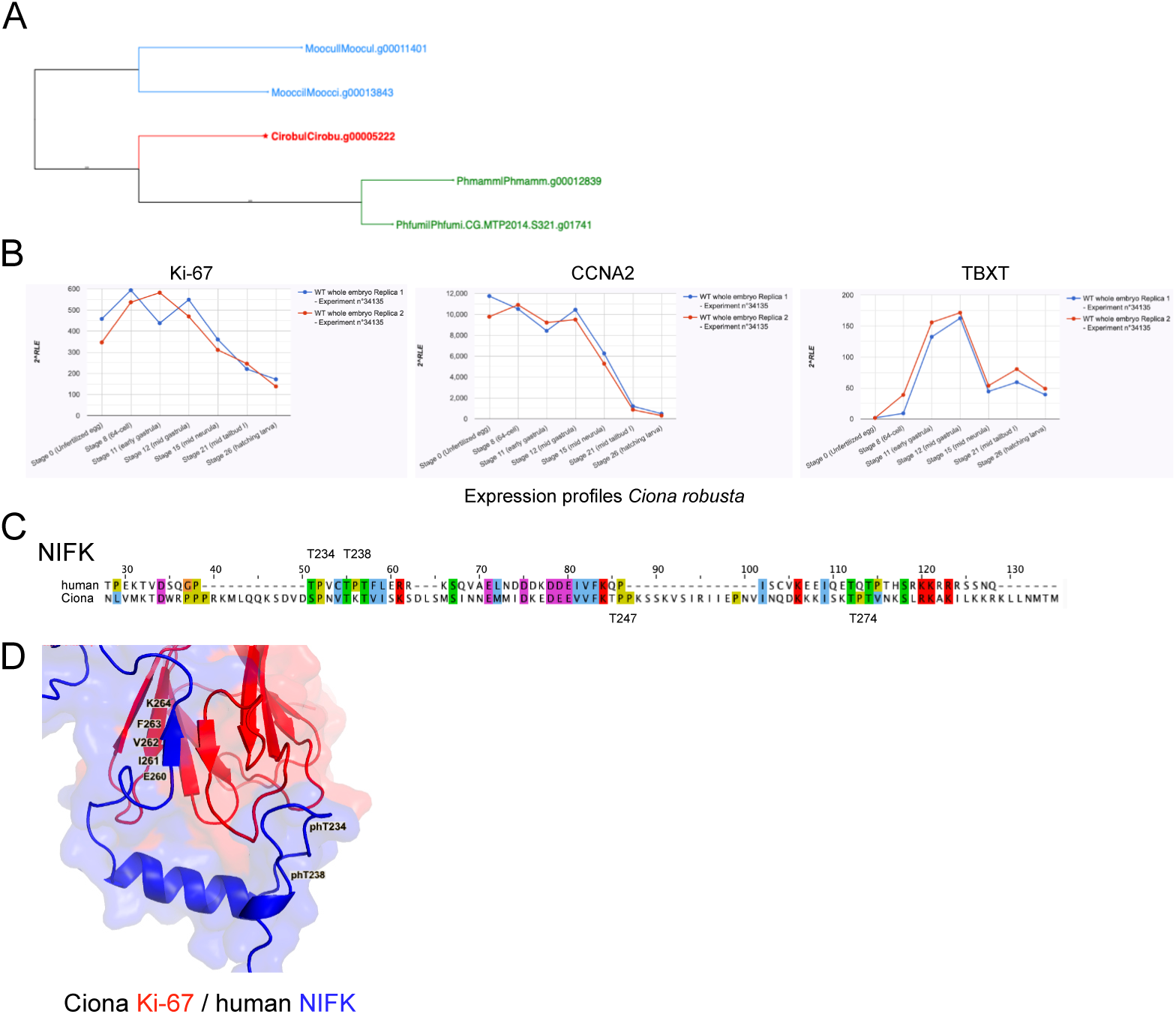
The expression profile, cellular localisation and interaction with NIFK are conserved between Ciona and huma Ki-67. A. Gene phylogeny of Ciona Ki-67 homologue (source: www.aniseed.fr). B. Expression profiles of Ciona Ki-67, Cyclin A and TBX transcription factor, from unfertilised egg to hatching larva (source: www.aniseed.fr). C. Sequence alignment of C-termina of human and Ciona NIFK; Thr residues that when phosphorylated interact with Ki-67 FHA domain are indicated (for Ciona, as predicted by AlphaFold3). D. AlphaFold3 model of Ciona Ki-67 FHA domain interaction with C-terminus of human NIFK phosphorylated on Thr234 and Thr238. The NIFK residues forming the antiparallel ß-strand and interaction of phThr are identical to that of human Ki-67-NIFK. Images generated using PyMol.

**Supplementary Figure 2.**
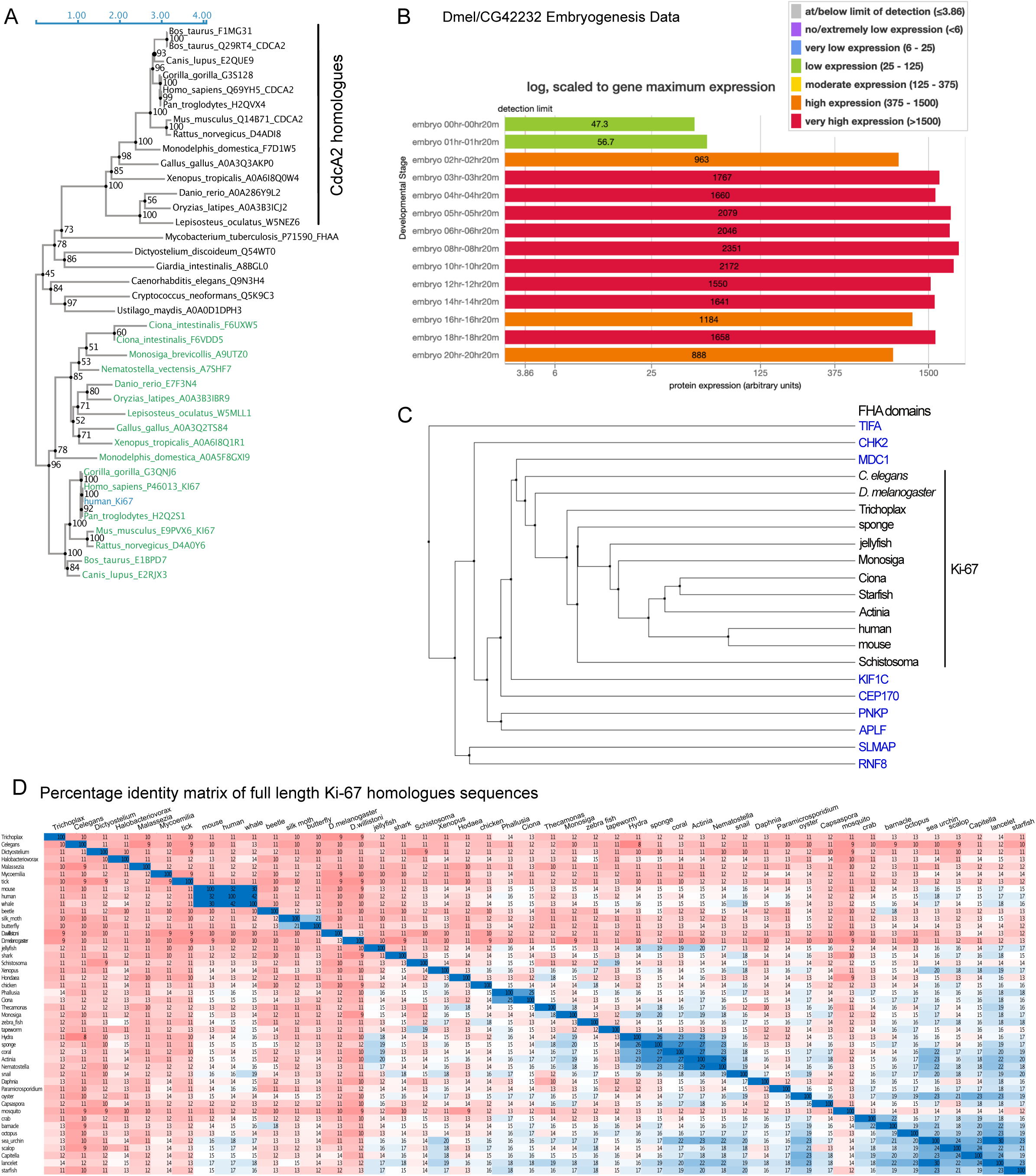
The *Drosophila* Ki-67 homologue is expressed in early embryonic stages and its overexpression in human cells induces chromatin compaction. A. Search for orthologues of human Ki-67 using the phylogenetic tool SHOOT (against all Metazoa) reveals various non-vertebrate species, including Ciona and *C. elegans* (Q9N3H4) homologues, as well as a separate branch of CDCA2 homologues. B. *Drosophila melanogaster* Ki-67 homologue (encoded by CG42232 gene; http://flybase.org/) expression profile in early embryogenesis. C. Phylogenetic tree reconstructed using BLOSUM62 substitution matrix following MUSCLE multiple sequence alignment of FHA domains of the indicated proteins. Human non-Ki-67 proteins are indicated in blue. D. Percentage identity matrix of full length Ki-67 homologues sequences (aligned by MUSCLE), presented in blue-to-red colour scale (red < 15%).

**Supplementary Figure 3.**
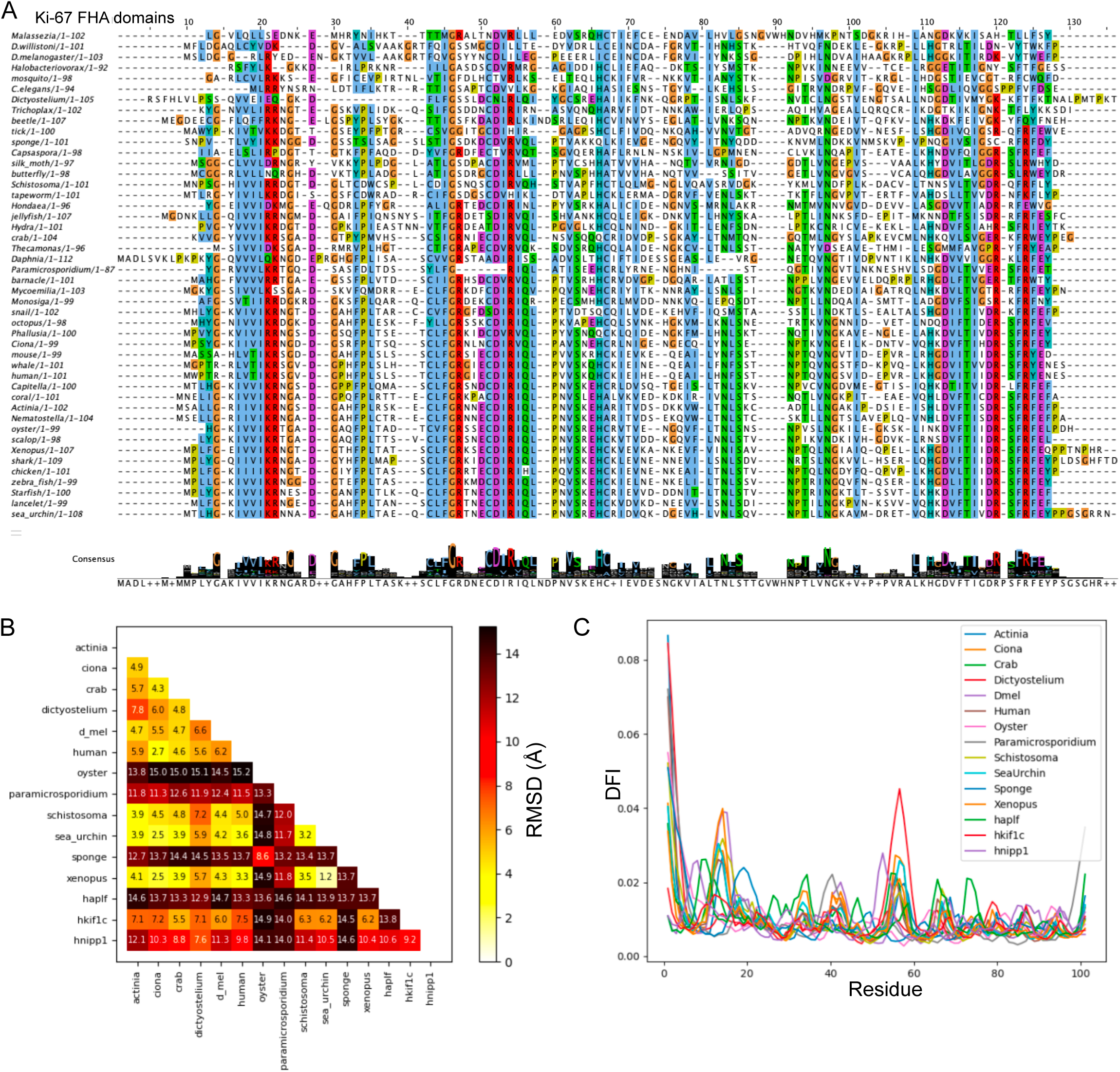
FHA sequences of Ki-67 homologues are highly conserved. A. MUSCLE sequence alignment of FHA domains of Ki-67 homologues (coloured by Clustal). B. Backbone RMSD (in °A) after 2.5 μs of MD. Representative structures of the dominant cluster from hierarchical agglomerative clustering were used for comparison. C. Raw DFI profiles for the FHA domains of Ki-67 homologues and of control human proteins (APLF, KIF1C, NIPP1).

**Supplementary Figure 4.**
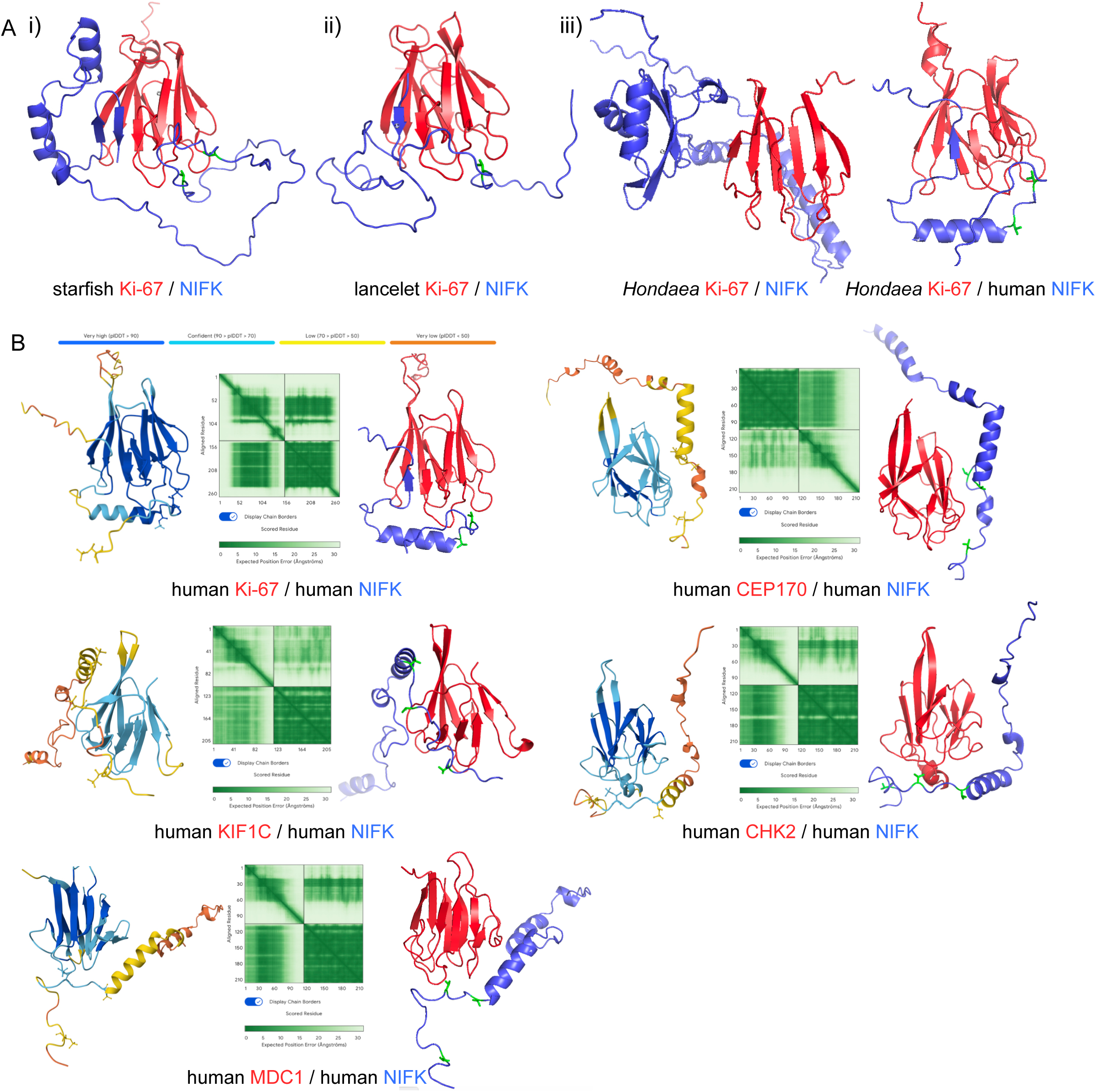
The interaction between Ki-67 FHA domain and phosphorylated C-terminus of NIFK is specific to Ki-67, with variations in different non-vertebrates. A. AlphaFold3 models of FHA domains of the indicated Ki-67 homologues with Thr-phosphorylated (indicated in green) NIFK C-terminal peptides, representing the different modes of interaction: i) FHA:NIFK interaction as in human (formation of the anti-parallel ß-strand by NIFK and N-terminal phThr binding to loops of FHA domain); ii) interaction FHA:NIFK similar to Ciona (formation of the anti-parallel ß-strand by NIFK, C-terminally of which phThr residues interact with FHA loops); iii) Ki-67 FHA does not interct with NIFK of the same species, but the interaction is conserved with human NIFK. Images were generated using PyMol. B. AlphaFold3 models of interaction of C-terminus of human NIFK (phThr234 and phThr238) with FHA domains of human Ki-67 (aa 1-110), CEP170 (aa 1-105), KIF1C (aa 498-597), CHK2 (aa 95-202) and MDC1 (aa 27-132). Each panel shows: left, the predicted structure coloured according to pLDDT (predicted local distance difference test), a per-atom confidence estimate on a 0-100 scale where a higher values indicate higher confidence (the colour code shown above); middle, PAE (predicted aligned error) matrix, an estimate of the error in the relative position and orientation between two tokens in the predicted structure, where higher values indicate higher predicted error and therefore lower confidence; right, predicted 3D structure generated using PyMol (NIFK in blue, phThr in green).

**Supplementary Figure 5.**
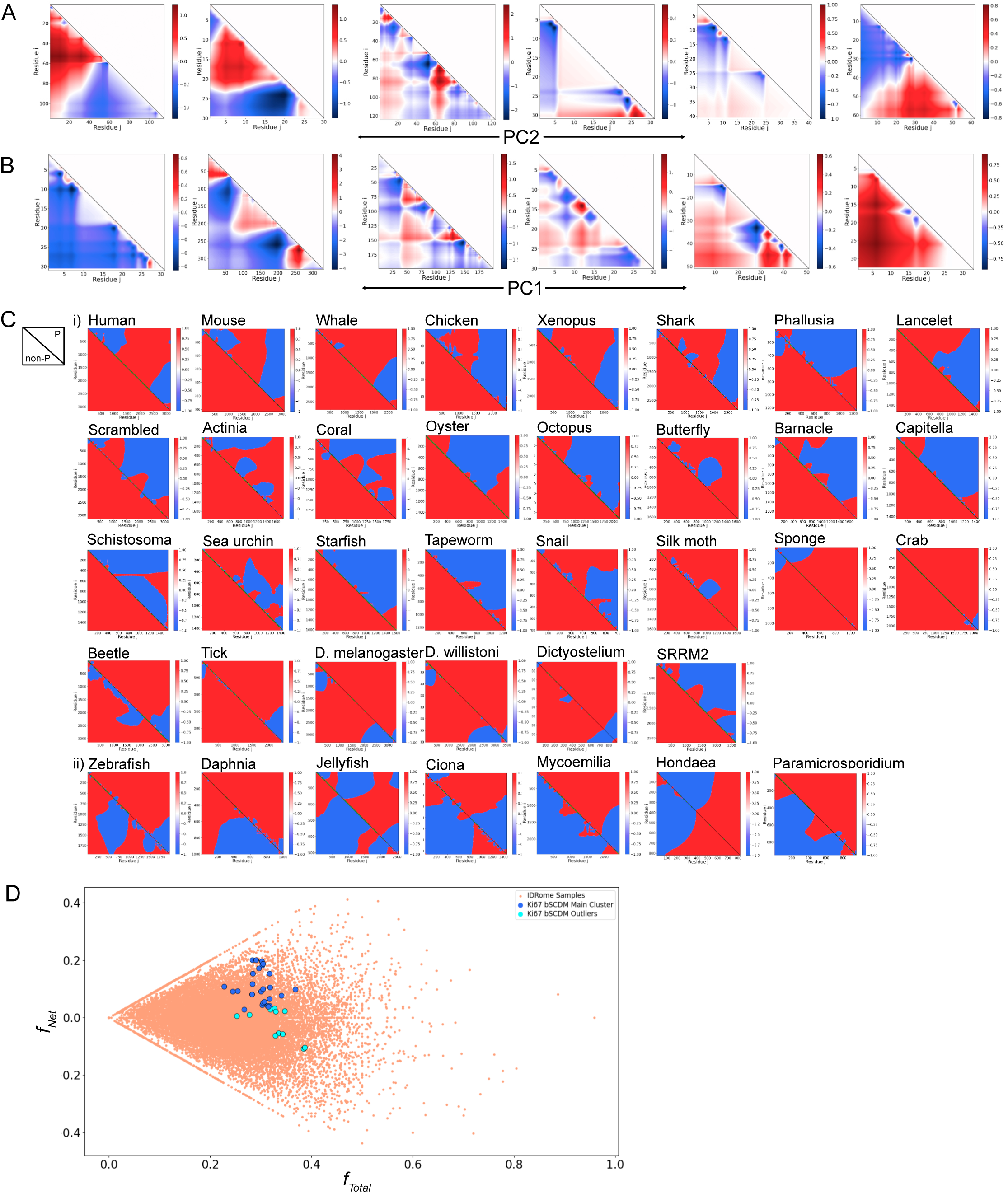
bSCDM profiles of Ki-67 homologues are distinct from the human IDRome bSCDM space. A. SCDM plots for IDRs along the PC1=0 line of Ki-67 bSCDM PC-space plot. Six representative IDRs were chosen along the PC1=0 line covering the whole range of PC2 values (-120 to 120). From left to right, PC2 values for IDRs are: -120.0, -69.9, -20.8, 30.1, 80.0, 119.7. B. SCDM plots for IDRs along the PC2=0 line of Ki-67 bSCDM PC-space plot. Six representative IDRs were chosen along the PC2=0 line based cevering the entire range of PC1 values (-130 to 120). From left to right, PC1 values for IDRs are: -130.0, -80.6, -28.7, 21.7, 68.0, 119.8. C. bSCDM matrices for Ki-67 homologue sequences. bSCDM graphs provide visual representation of single chain electrostatic interactions (unmodified sequences in the bottom triangle, and phosphorylated sequence in the top triangle). Two groups of sequences were identified: i) uniform repulsive; and ii) mixed repulsive / attractive when unphosphorylated. D. Fraction of net charge (net charge per residue, ƒ_+_ - ƒ-) vs fraction total charge (fraction of charged residues, ƒ_+_ + ƒ-) for Ki-67 proteins and the IDRome. IDRome proteins are colored in salmon, Ki-67 proteins part of the central cluster in bSCDM PC-Space are colored in blue, and the outliers are colored in light blue.

**Supplementary Figure 6.**
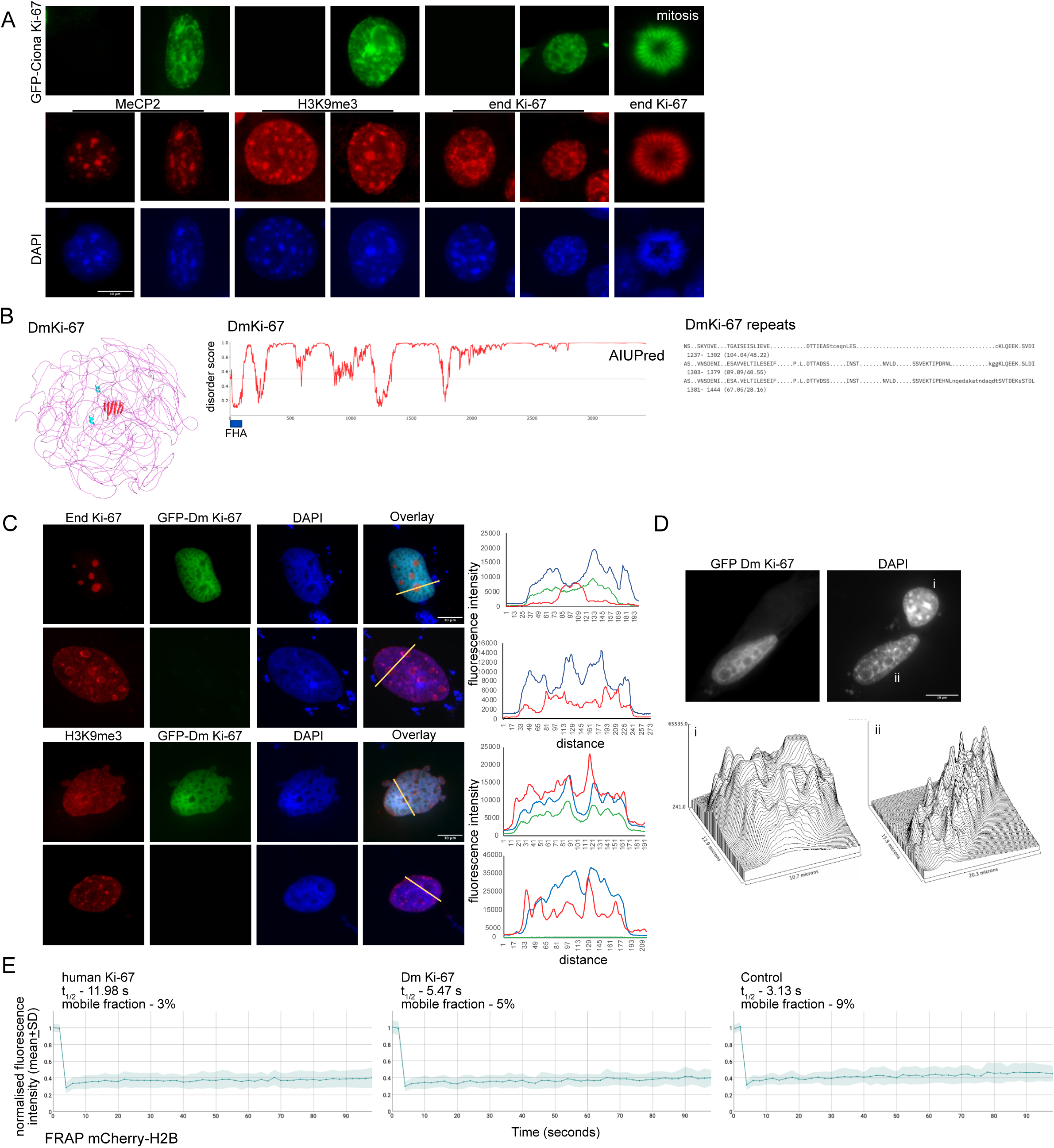
Ciona and *Drosophila* Ki-67 homologues regulate chromatin organisation without altering histone dynamics. A. Immunofluorescence images of mouse 4T1 cells overexpressing GFP-Ciona Ki-67, stained with the indicated antibodies; DNA was stained with DAPI. All cells are shown at the same magnification; scale bar 10 µm. B. Left, AlphaFold3 model of *Drosophila melanogaster* Ki-67 (FHA domain in red). Middle, AIUpred disorder prediction plot of the DmKi-67 homologue; N-terminal FHA domain is indicated. Right, repeats identified in the DmKi-67 IDR by RADAR. C. HeLa cells overexpressing GFP-*Drosophila* Ki-67, stained with the indicated antibodies; DNA was stained with DAPI. All cells are shown at the same magnification; scale bar 10 µm. Right, fluorescence intensity profiles through the region indicated on the overlay image. D. Top, immunofluorescence images of mouse NIH3T3 cells overexpressing GFP-DmKi-67; DNA was stained with DAPI; scale bar 10 µm. Bottom, surface immunofluorescence intensity plots for the untransfected (i) and DmKi-67-expressing (ii) cells. E. FRAP analysis of mCherry H2B in control untransfected, GFP-human Ki-67, and GFP-*Drosophila* Ki-67-transfected cells. Mean normalised fluorescent intensity (+ SD) recovery curves are shown.

**Supplementary Figure 7.**
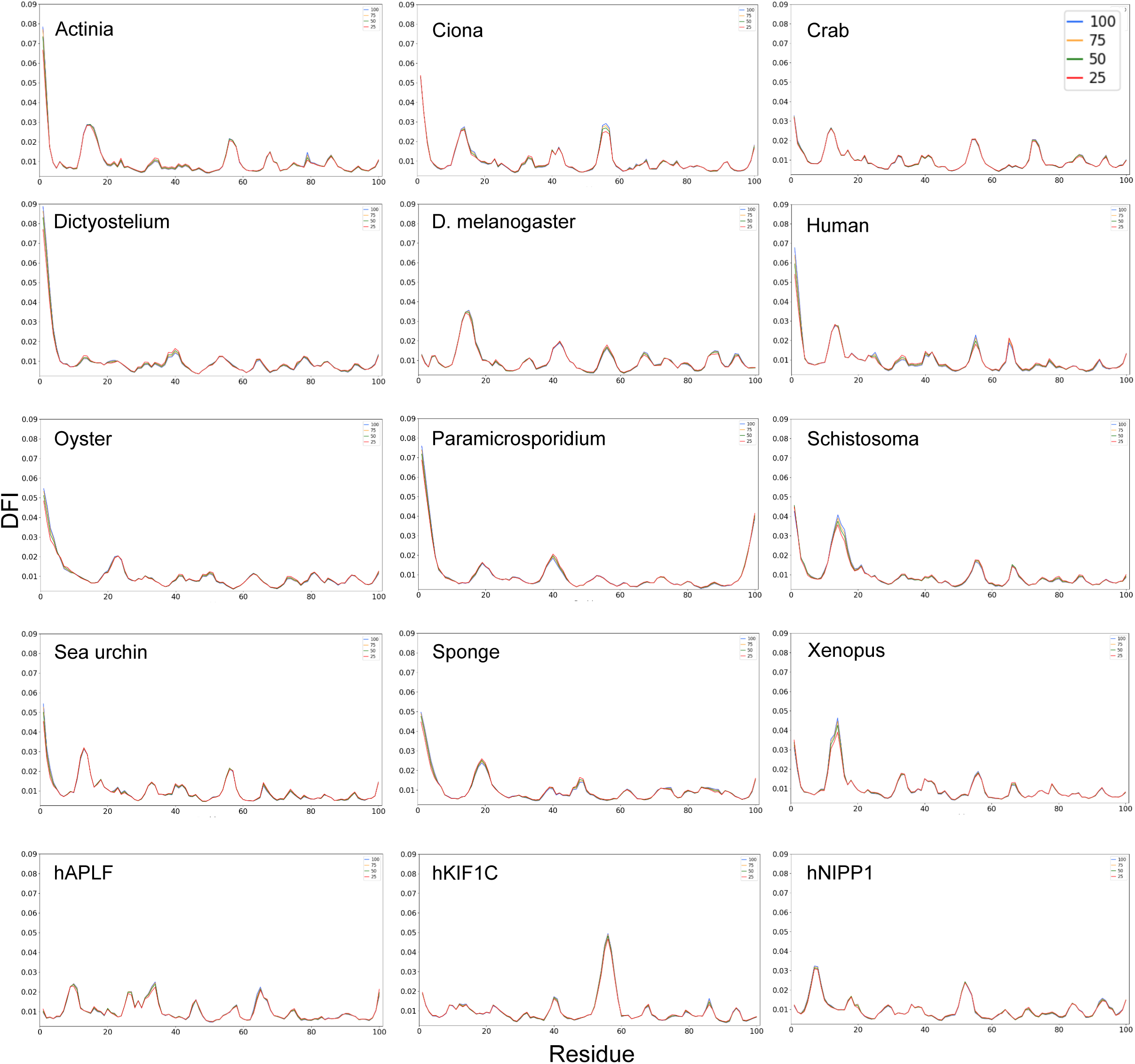
Raw DFI profiles of FHA proteins with varying window sizes: 25ns, 50ns, 75ns, and 100ns. These similarities along with self consistency checks performed on the trajectories themselves ensure robustness of the results.

